# Stay clear and dry! How microstructure diversity can offset the hydrophobicity costs of transparency in clearwing Lepidoptera

**DOI:** 10.1101/2021.10.11.463886

**Authors:** Doris Gomez, Jonathan Pairraire, Charline Pinna, Monica Arias, Céline Houssin, Jérôme Barbut, Serge Berthier, Christine Andraud, Thierry Ondarçuhu, Marianne Elias

**Author notes:** **CORRESPONDING AUTHOR:** Doris Gomez.

## Abstract

Living organisms are submitted to multiple developmental and selective constraints resulting in evolutionary compromises, one of the best examples being the integument (the outer protective layer of living organisms) which is fundamentally multifunctional. Integument anti-wetting or hydrophobicity – evolved in relation to complex and various structures - is a crucial property as it serves multiple functions like self-cleaning, locomotion, or defence against pathogens and may interfere with other functions like thermoregulation or communication. Elucidating the structure-property relationships and unravelling potential trade-offs is crucial to understand the evolution of the integument. In opaque Lepidoptera, wing scales actively contribute to anti-wetting. In clearwing Lepidoptera, wing scales are often reduced, raising the question of whether they can maintain similar hydrophobicity levels to those of opaque species and if not, whether wing microstructure (scale density, shape, insertion, and coloration) may mitigate the costs of a lower hydrophobicity. To answer these questions, we measure static contact angle (CA) of water droplets at different stages of evaporation in opaque and transparent patches of 23 Lepidoptera species that show a high diversity in wing microstructure. More specifically, we find that transparency is costly for hydrophobicity, and that such cost depends on wing microstructure. In general, transparent patches lose more hydrophobicity with water evaporation than opaque patches. Yet, this loss of hydrophobicity is attenuated for higher scale densities, erect scales compared to flat scales, coloured scales (for erect scales), multiple scale layers (for flat scales), or when combining two types of scales (piliform and lamellar) than having only one type of scale (piliform or lamellar). Nude membranes show the lowest hydrophobicity values. We find that wing hydrophobicity negatively relates to optical transparency, showing a trade-off between optics and hydrophobicity. Moreover, we find that tropical species have higher hydrophobicity in their transparent patches than temperate ones, suggesting transparent patches are under stronger selection for hydrophobicity in tropical than in temperate species. These novel findings, which are consistent with the physics of hydrophobicity, suggest that insect wings are evolutionary multifunctional compromises.

## INTRODUCTION

Living organisms are submitted to multiple developmental and selective constraints resulting in evolutionary compromises. The integument, the outer protective layer of plants and animals, perfectly illustrates this concept as it is involved in multiple functions like mobility, thermoregulation, communication or camouflage, and defense against pathogens. Hydrophobicity is a crucial property of the integument in terrestrial organisms: avoiding unwanted water efficiently contributes to self-cleaning mechanisms and defense against pathogens, as water droplets that roll off remove potentially contaminating particles, like dust or bacteria (Wagner et al., 1996). Hydrophobic surfaces, by their structure, can have direct bactericidal effects, such as in geckos (Watson et al., 2015) and cicadas (Ivanova et al., 2012). Anti-wetting also contributes to locomotion: in geckos, the hydrophobicity of complex toepads allows adhesion and locomotion on any surface (Autumn et al., 2014); in flying insects, the hydrophobicity of wings – which reduces drag, removes weight, and limits wing damage – enhances flight ability, and extends it to rainy conditions (Watson et al., 2011). In water striders, the hydrophobicity of legs ensure water skating (Gao & Jiang, 2004) and the hydrophobicity of the body cuticle creates a plastron (air-film) which helps under-water respiration, provides buoyancy, and protects from submergence, thereby facilitating the colonization of the open ocean (Mahadik et al., 2020). Yet, hydrophobicity has its limits: for instance, water films, which help cooling through water evaporation, cannot form on hydrophobic surface; hence hydrophobicity can interfere with thermoregulation potentially limiting cooling mechanisms and efficiency. There can exist trade-offs between antagonistic needs and different evolutionary compromises can be selected in various environmental conditions. Finally, the hydrophobicity property has evolved in relation to integument structure: in geckos, hydrophobic toepads have been gained and lost multiple times independently, leading to a structural diversity at all scales (Gamble et al., 2012), which relates to differences in locomotion performance and habitat use between species (Elstrott & Irschick, 2004). Hence, deciphering the structure-property variations and unraveling potential trade-offs is crucial to understand the evolution of plants and animals.

As predicted by physics (Wenzel, 1936) and illustrated in plants (Barthlott & Neinhuis, 1997), a key parameter for hydrophobicity is surface texture or roughness. A water droplet on a textured hydrophobic surface can exhibit two different wetting states. In the Cassie-Baxter state (Figure 1, series a), the water droplet sits on top of the texture, with trapped air underneath and cavities filled with air (composite state, solid and air in contact with water under the drop), and hydrophobicity is at a maximum. If this state is thermodynamically metastable, the water droplet may undergo the so-called Cassie-Baxter to Wenzel transition, in which water penetrates the air-filled cavities by capillarity. In the Wenzel wetting state (Figure 1 series b), the water droplet fully fills all the cavities of the texture and adheres to the surface (non-composite state, only solid in contact with water under the drop), decreasing the surface energy; hydrophobicity is then lost (Hasan et al., 2012). Compared to the Wenzel state, the Cassie-Baxter state is of high biological interest as it offers an incomplete water-surface contact and a weak water adhesion. Maintaining a stable Cassie-Baxter state is crucial to maintain high hydrophobicity under harsh environmental conditions, like rainfall.

**Figure 1.**
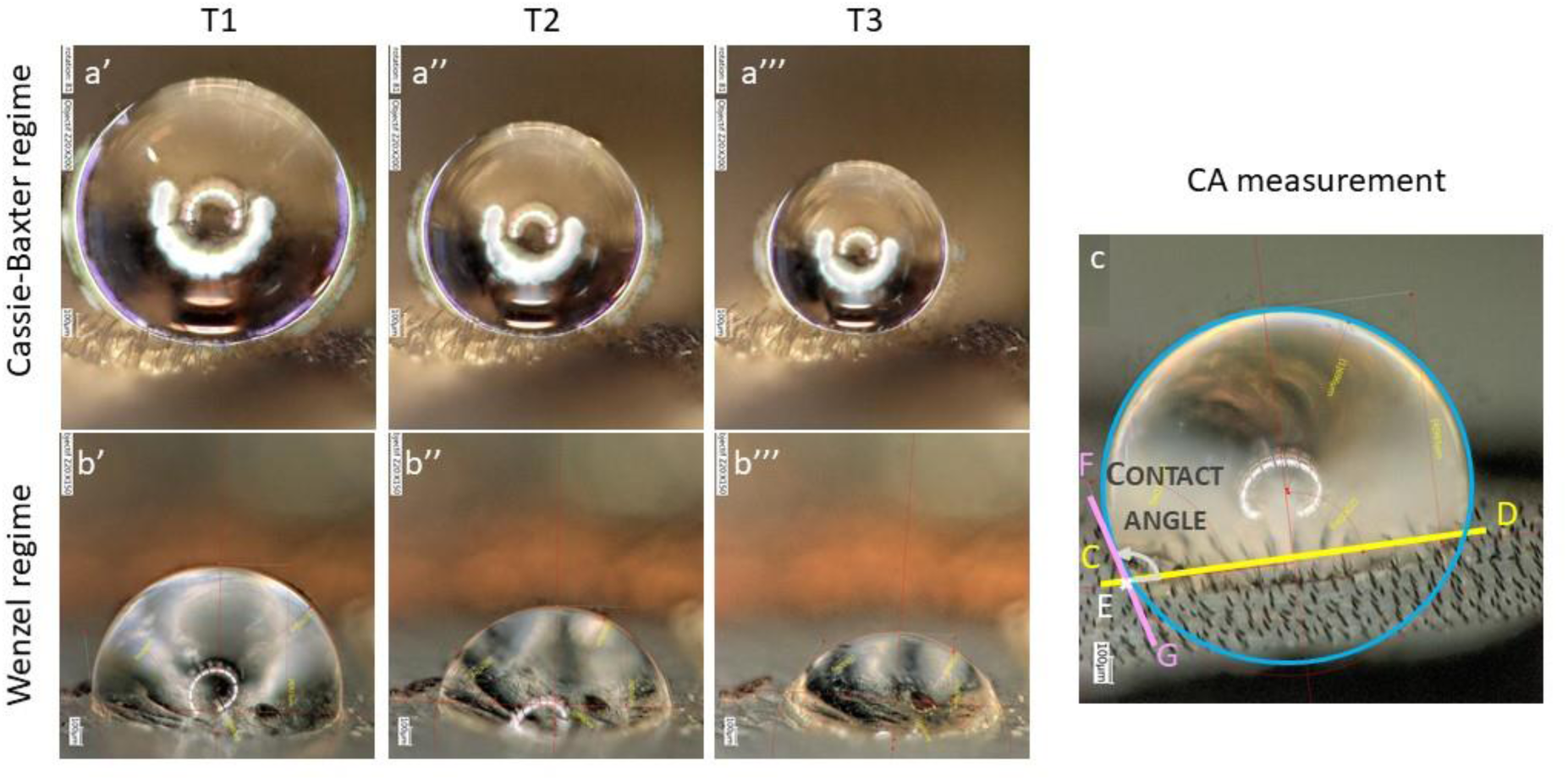
Examples of water droplets dropped in the transparent zone: Cassie-Baxter regime (series a) for *Eutresis hypereia* combining erected coloured piliform and lamellar scales and Wenzel regime (series b) for *Neorcarnegia basirei* with a nude membrane. Water droplet evolution is shown at different times: T1 (a’, b’), T2 (a’’, b’’), and T3 (a’’’, b’’’). Principle of contact angle (here called CA) measurement on a photo example from *Parantica sita* (c): first, we draw a theoretical circle on the droplet shape (blue). We then figure the wing surface (yellow segment CD). We define the point E (white) as the intersection of the water droplet and the segment CD and the segment FG (pink) as the tangent to the circle in E. We then compute the contact angle expressed in degrees as the angle between (CD) and the tangent (FG) of the water droplet at the point E. The rounder the droplet, the higher the CA value.

Roughness at nanoscale – the parameter most studied to date in animals and plants - increases hydrophobicity, as shown in cicadids and dragonflies (Byun et al., 2009; Oh et al., 2017; M. Sun et al., 2009). Yet, multiscale roughness –at nano and micro scale – is even more efficient: it increases hydrophobicity and its thermodynamic stability and it reduces water adhesion. This was shown in modelling studies (Bell et al., 2015; Su et al., 2010) and repeatedly illustrated in various biological examples: in the Lotus (so-called ‘Lotus effect’) and other plants (Barthlott et al., 2016) as well as in insects such as water striders (Gao & Jiang, 2004) and mosquitoes (Wu et al., 2007). Increasing thermodynamic stability allows maintaining hydrophobicity with water droplets of various sizes (dew, fog, rain) and increases anti-fogging properties, i. e. the resistance to tiny water droplets condensing on the surface. While the role of nanostructures in hydrophobicity has been extensively documented (e.g. Patankar, 2004; Porcheron & Monson, 2006), the role of microstructures shape in determining hydrophobic properties has been limited to simple geometries (cylinders in Cansoy et al., 2011; cones in Ding et al., 2019; P. Tsai et al., 2010) and remains poorly investigated from an empirical perspective. The only existing empirical studies with such an approach either describe the variations in hydrophobicity between various micro-architectures but without invoking explanations (Sanchez-Monge et al., 2015) or focus on one type of micro-architecture (Fang et al., 2015; Oh et al., 2017; G. Sun & Fang, 2015), inescapably showing a major influence of nanostructures to explain the variations in hydrophobicity, since nanostructures are the main elements varying among these species.

Lepidoptera (from the ancient Greek λεπίς: scale and πτερόν: wing) – butterflies and moths – offer an outstanding study system to investigate this question. They are typically characterized by large wings entirely covered with flat and coloured lamellar microscopic scales (Ghiradella, 1998; Miaoulis & Heilman, 1998). Scales are in average around 100 µm long and 50µm wide. Through their pigmentation and structure, scales are involved in multiple functions such as antipredator defences (e. g. camouflage, deflection, mimicry in Cuthill et al., 2005; Stevens et al., 2008), communication (Kemp, 2007), thermoregulation (Berthier, 2005; Krishna et al., 2020; Miaoulis & Heilman, 1998; C.-C. Tsai et al., 2020) and flight enhancement (Nachtigall, 1967; Slegers et al., 2017). They also confer superhydrophobic properties to the wing, resulting in water repellency and self-cleaning (Wagner et al., 1996; Wanasekara & Chalivendra, 2011). Superhydrophobicity *sensu lato* is defined by water droplets making high contact angles (>150°) with a surface. Self-cleaning – superhydrophobicity *stricto sensu* (definition not taken here) – adds to this condition a weak water adhesion, estimated by a minimal tilt from the horizontal plane needed for water droplets to roll-off (roll-off angle of a few degrees) or a minimal hysteresis (difference between advancing and receding contact angles). Superhydrophobicity is thus a *sine qua non* condition for water repellency and self-cleaning. Opaque butterflies and moths typically have self-cleaning wings, as attested by small roll-off angles (Fang et al., 2015, 2017). Scarce relevant studies suggest that wing hydrophobicity may depend on wing microstructure (scale presence in Finet et al., 2023; scale type and insertion angle in Perez Goodwyn et al., 2009; presence and type of scale in Wagner et al., 1996), and on wing macrostructure: species with longer wings (Byun et al., 2009), or larger ratio of wing area to body mass (Wagner et al., 1996) show higher hydrophobicity and wing shape was invoked to explain natural variations in hydrophobicity (Byun et al., 2009).

While the vast majority of Lepidoptera species has opaque wings, some species from various lineages show transparent or translucent wings (Gomez et al., 2021). While the evolution of transparency in an order of insects that typically harbour large wings covered by coloured scales may appear puzzling, recent experiments have shown that transparency is beneficial to butterflies and moths because it reduces their detectability from visually-hunting predators (Arias, Elias, et al., 2020; Arias et al., 2019; McClure et al., 2019). In Lepidoptera, wing transparency is involved in various anti-predator defences, spanning from camouflage (Arias, Elias, et al., 2020) to Batesian mimicry with Hymenoptera (Skowron Volponi et al., 2018) and masquerade (Arias, Barbut, et al., 2020; Costello et al., 2013). Transparency is associated with a broad microstructural diversity (i. e. scale diversity, see examples in ESM, Figure S1), the membrane being nude or covered with scales varying in type (piliform, i. e., hair-like, and/or lamellar), insertion on the membrane (flat or erect), and colouration (coloured or transparent) (Gomez et al., 2021). All combinations of scale type, insertion, and colouration (i. e., structural strategies Gomez et al., 2021) can be found in nature (ESM, Figure S1), and they differ in their efficiency at transmitting light : the nude membrane are most efficient while flat coloured scales (lamellar alone or in combination with piliform) are least efficient (Gomez et al., 2021). Microstructures are complemented by nanostructures on the scales and on the wing membrane. Membrane nanostructures reduce reflection levels and increase light transmission (Pinna et al., 2021; Pomerantz et al., 2021; Siddique et al., 2015; Yoshida et al., 1997).

Because transparency often entails profound modifications – and, in the vast majority of cases, reduction -of scale dimensions and scale density (Gomez et al., 2021), it can be hypothesized that achieving optical transparency may come at a cost to hydrophobic performance, both in water repellency and self-cleaning properties. This cost manifests as a trade-off which arises when two functional requirements cannot be simultaneously optimized, since structural features that enhance one property may inherently compromise the other. Water repellency and self-cleaning are vital for butterflies and moths, which are large-winged insects: water repellency is crucial for flight and for preventing wings from sticking together, especially in tropical rainforest species with daily rain and high humidity. Likewise, self-cleaning helps removing dust contamination that impairs flight (Wagner et al., 1996). Among the lepidopteran species investigated so far for hydrophobicity (Fang et al., 2015; Perez Goodwyn et al., 2009; Wagner et al., 1996; Wanasekara & Chalivendra, 2011; Zheng et al., 2007), only four clearwing butterfly species have been included: *Parantica sita* (with lamellar titled scales) and *Parnassius glacialis* (with flat lamellar scales), with high or moderate hydrophobicity respectively (Fang et al., 2015; Perez Goodwyn et al., 2009), *Phanus vitreus* (with erect lamellar scales) with lower hydrophobicity in transparent than opaque areas (Finet et al., 2023), *Greta oto* (with piliform scales) with one of the lowest hydrophobicity values found in butterflies (Wanasekara & Chalivendra, 2011). Scarce data suggest that a greater reduction in scale dimensions or coverage on the wing membrane may entail higher costs in terms of hydrophobicity, as suggested by the lower hydrophobicity in transparent areas of *Phanus vitreu*s with removed scales than with scales (Finet et al., 2023). However, large-scale comparative studies are currently lacking.

To fill that knowledge gap, we here explore to what extent anti-wetting ability is influenced by scale microstructure in species, whether it entails a trade-off with optical transparency and whether it depends on climatic conditions, by selecting a subset of 23 species (ESM, Figure S2) from a broad study of 123 clearwing Lepidoptera species (Gomez et al., 2021) that show a large diversity in microstructure. In these species, we explored the links between microstructure, macrostructure, hydrophobicity and optics while controlling for phylogenetic relatedness between species. We measured the contact angle (CA) made by water droplets on the wing at different stages of water evaporation (thus droplet size). With these measurements, (i) we explored the relationships between hydrophobicity and wing macrostructure. (ii) We then explored the relationships between hydrophobicity and wing microstructure, i. e. structural strategy (presence or absence of scale, scale shape, scale insertion angle, coloration and density). (iii) To go beyond the associations beyond hydrophobicity and structural strategy, we explored the geometry of some structural strategies of particular interest (erect versus flat geometries, geometries involving two types of scales),. We tested whether there existed consistent associations between geometrical scale features (scale dimensions, spacing, density) within the structural strategies that could explain the observed variations in hydrophobicity. (iv) To identify the selective pressures acting on hydrophobicity, we tested whether hydrophobicity and light transmission showed potential trade-off or synergy. A trade-off between hydrophobicity and light transmission would reveal a cost of transparency for water repellency. If microstructures play a dominant role in conferring hydrophobicity, species most efficient at transmitting light – which lack scales or have highly modified scales, resulting in low coverage of the wing surface – are expected be less efficient at repelling water. (v) Finally, to identify whether hydrophobicity is influenced by climatic conditions, we tested the links between the latitude of species habitat and hydrophobicity: if repelling water is more important in the tropics where rain and humidity are inescapable, tropical species are expected to show higher hydrophobicity than temperate species.

## METHODS

### Species selection

Scale type and scale insertion have been suggested to influence hydrophobicity (Perez Goodwyn et al., 2009). Scale coloration, often involving melanin deposition which increases cuticle hardening in insects (Sugumaran, 2009), could increase scale stiffness and ability to repel water droplets. Hence, we selected a set of species varying in structural strategies – scale type (N=nude membrane, P=piliform bifid or monofid scales, L=shape different than piliform, hereafter called lamellar, or PL=association of piliform and lamellar scales), insertion (E=erect or F=flat), and colouration (C=coloured or T=transparent) – from the study of 123 species of clearwing Lepidoptera (Gomez et al., 2021). We minimized the phylogenetic relatedness between species harbouring the same type of structural strategies to increase the power of comparative analyses. We selected a total of 23 species from 10 families (Figure 1 & ESM, Figure S1, list in ESM Table S1), comprising 3 species for the structural strategies (N, PFC, PEC, LFC, LFT, LET), 2 species for PLEC and LEC, and 1 species for PLET, as for some species only a limited number of specimens were present in the collections. For each species, we selected three specimens in good condition either from Paris MNHN collections or from our own private collections. 54/69 specimens (all species but *Eutresis hypereia*) had labels with exact collect location that could be tracked down to GPS coordinates.

### Hydrophobicity measurements

We measured the static contact angle of water droplets and wing surface in the transparent and opaque zones of the forewing of three museum specimens per species, and we monitored contact angle at three times, as water evaporated and droplet size decreased (Figure 1). For each specimen, we used a purpose-built water-droplet dispenser (a graduated pipette on a holder) and a Keyence VHX-5000 microscope (equipped with Z20 zoom) to image water droplets on butterfly wings. As a general procedure, we dropped a series of three 1 µl water droplets (volume usually taken to assess hydrophobicity (Hasan et al., 2012; Perez Goodwyn et al., 2009)) at three locations of the transparent and opaque zones of the dorsal side of a wing. After the water droplet was dropped (time T1), we allowed its volume to be approximately divided by two (time T2) and by four (time T3) compared to its original volume. Since evaporation kinetics depended on droplet shape, time intervals elapsed between consecutive photos were not identical from one species to another. At each time, we took a photo in which we measured the static contact angle (measurement principle and examples in Figure 1). The contact angle measured at time T1 can be considered as the advancing contact angle and thus serves as an indicator of surface hydrophobicity. The subsequent decrease in CA due to water evaporation, hereafter referred to as the “loss of hydrophobicity,” reflects contact angle hysteresis and is indicative of the surface’s self-cleaning ability. A smaller loss of hydrophobicity corresponds to a reduced contact angle of hysteresis, and an improved self-cleaning performance.

Our protocol only included the measurement of static contact angles which can potentially vary within a range of possible metastable values (Liu et al., 2019). Yet, statistical analyses showed that contact angle values were highly repeatable (i) for the same wing, zone and time, (ii) for both wings in the same zone, and (iii) for the same species. This ensured our protocol yielded reliable values (see detailed methods and results in ESM Table S2). We thus kept the same protocol, but we measured only the forewing.

We did all measurements on dry museum specimens, as widely done in comparative studies of hydrophobicity (Fang et al., 2015; G. Sun & Fang, 2015; Wagner et al., 1996; Wanasekara & Chalivendra, 2011). Desiccation makes wings flatter and more comparable but it may alter the relative hydrophobic behaviour of the different species, i.e. the more/less hydrophobic species in dry conditions may not be the more/less hydrophobic species in humid conditions. In a restricted sample of 5 species showing the most common microstructures (see ESM for details), we measured the contact angle in the opaque zone and the transparent zone of a dry specimen and again on the same points, once the specimen rehydrated for 48h and showed that species ranking was conserved between dry and humid treatment, be it in the opaque or in the transparent zone, confirming the validity of our protocol (see ESM for details).

### Measurements of wing macrostructure and microstructure

To characterize wing macrostructure, we took photos of the three specimens of each species using a camera (D800E Nikon, 60mm lens, annular light). We analysed photos using ImageJ (Schneider et al., 2012). Given the role of wing length (Byun et al., 2009), wing shape (Byun et al., 2009), and ratio of total wing area to body mass (Watson et al., 2008) on hydrophobicity and self-cleaning ability, we computed wing length, length-to-width LW ratio and the ratio of total wing area to body volume, taking the volume as a proxy for mass for dry specimens, and assuming the body to be a cylinder, for which we measured length (thorax+abdomen) and width. Using the ‘rptR’ package (Stoffel et al., 2017), we found that all wing macrostructural measurements were repeatable, i.e. that a specimen was representative of its species for all wing macrostructural variables (ESM Table S2).

To characterize wing microstructure (i.e. scale characteristics, presence, type, insertion, coloration, density), we imaged the dorsal side of forewing transparent and opaque zones using microscopes (Zeiss Stereo Discovery V20 and Keyence VHX-5000). We did that in one specimen per species because scale dimensions and density were repeatable at zone by species level in Gomez et al. (2021). Using ImageJ and Keyence built-in tool, we measured scale density (per mm²), length and width (µm), scale surface (in µm²) as the product of length by width, and scale coverage as the product of scale surface (expressed in mm²) by scale density. We counted the number of different scale types: 0=nude membrane, 1= lamellar or piliform, 2= lamellar and piliform. For flat lamellar scales, we also computed the density of scale top layer and computed the number of layers as the ratio between density and top layer density.

Not only the presence of multiscale roughness is important for hydrophobicity, but its spatial geometry is crucial for its stability (various fractal geometries tested in Bittoun & Marmur, 2012). For scale geometry (presence of one type of scales, either piliform or lamellar scales but not both), we investigated the variations in scale length or scale width with scale insertion on the membrane as this was anticipated to greatly influence the spatial geometry of wing surface, and with scale coloration as melanin was a component of cuticle hardening in insects (Sugumaran, 2009). From Gomez et al.’s broad study (2021), we selected the 96 species with one type of scales only. When both piliform and lamellar scales were present, we investigated how these two types of scales were spatially associated as it was potentially the most complex micro-architecture. From Gomez et al.’s broad study (2021), we selected the 8 species that had a PL strategy, i.e. a combination of piliform scales and lamellar scales in the transparent zone. These species were *Athesis clearista, Diaphania unionalis, Dysschema boisduvalii, Eutresis hypereia, Macrosoma conifera, Methona curvifascia, Nagara vitrea, Praeamastus fulvizonata.* In these species, we computed (i) the ratio in length between piliform scales and lamellar scales, (ii) the ratio in density between piliform scales and lamellar scales, and (iii) the spatial association between piliform scales and lamellar scales.

### Optical measurements

For one specimen per species, we measured specular transmittance from 300 to 700 nm as in Gomez et al. (2021), using a deuterium-halogen lamp (Avalight DHS), direct optic fibres (FC-UV200-2-1.5 × 100) and a spectrometer (Avaspec-2048 L, Avantes). Wing samples were placed perpendicular at equal distance between fibres aligned 5 mm apart (1 mm diameter spot). We took five measurements of the forewing in various points of the transparent zone. We computed the mean transmittance over [300-700] nm, which described the level of optical transparency. Optical measurements had been found highly repeatable at species level (ESM Table S2, and Gomez et al., 2021).

### Comparative analyses

To explore the questions outlines below, we ran Bayesian mixed models with Markov Chain Monte Carlo, correcting for phylogenetic relatedness, using the R package MCMCglmm (Hadfield, 2010) and the maximum clade credibility (MCC) phylogeny obtained in Gomez et al. (2021) and pruned to targeted species. According to the analysis, we pruned it to 23 species (for CA analyses, Figure S2), to 96 species (for P or L geometries) or to 8 species (for PL geometries). We tested different random factors (phylogeny, species, specimen) and retained the random assemblage that minimized DIC or if giving similar DIC values, the simplest in structure. Chain convergence was assessed visually and with Heidelberg’s and Geweke’s convergence and stationarity diagnostic functions from the R package coda (Plummer et al., 2006). We adjusted the number of iterations, the burn-in and the thinning to ensure best convergence and stationarity diagnostic and an effective sample size for all fixed and random parameters of at least 1000. Models were run with uninformative prior for random effect and residual variances (V = 1, nu = 0.002) not to constrain the exploration of parameter values. We selected the best model with a backward selection of fixed parameters based on Bayesian P-value important (95% credibility intervals excluding zero) or less important (90% credibility interval excluding zero).

With Bayesian models, we analysed the variation in contact angle with (i) wing macrostructure descriptors – time, zone, forewing length, surface, LW ratio, the ratio of total wing area divided by body volume, and relevant interactions –; (ii) wing microstructure descriptors – time, wing length (to correct for variation in scale dimensions), scale length, width, density, scale type, number of different types, scale insertion, scale colouration, and the number of layers. (iii) we tested whether in some structural strategies of interest (erect versus flat geometries, involving one or two types of scales), there existed consistent associations between scale geometrical features (scale length and width, scale spacing and density) to quantify the geometrical bases of variations in hydrophobicity. For instance, in structural strategies based on both scale types (piliform and lamellar), we analyzed length ratio, density ratio and spatial association between the two scale types in relation to scale insertion (see ESM for details). (iv) We tested for a potential trade-off between optical transparency and wing hydrophobicity, considering all measurements of contact angle, individual mean values, or species mean values at T1. (v) Finally, we tested whether tropical species were more hydrophobic than temperate species. To do so, we related for each specimen its average CA value to its latitude to the equator, the proportion of wing area occupied by transparency and wing length, while taking species as random effect, for the opaque and transparent zone separately. Gomez et al (2021) found a positive relationship between optical transparency and wing surface area covered by transparency. Hence, finding a relationship between CA and optical transparency could be simply explained by latitudinal variations in the proportion of wing surface area covered by transparency, which we also tested.

## RESULTS

### Variation in hydrophobicity and relation to wing macrostructure

We observe a general decrease in hydrophobicity with water evaporation in the opaque zone and in the transparent zone (Figure 2). Transparency appears costly: (i) the transparent zone shows lower hydrophobicity than the opaque zone of the same wing, whatever the size of the water droplet considered (zone effect in Figure 2A and in ESM Table S3, see ESM Figure S4 for distribution of hydrophobicity levels). In addition, (ii) we observe a stronger decrease in hydrophobicity with water evaporation in the transparent zone compared to the opaque zone of the same wing (time x zone effect in Figure 2A, and ESM Table S3). CA does not correlate to wing length, to the wing area to body volume ratio, or to the elongation of the forewing (ESM Table S3). Yet, species with more elongated wings, or with smaller wing area relative to their body or with shorter wings exhibit a greater loss of hydrophobicity with evaporation (negative forewing LW ratio x time, positive Wing Area to Body Volume x time, and positive wing length x time interaction effects in ESM Table S3).

**Figure 2.**
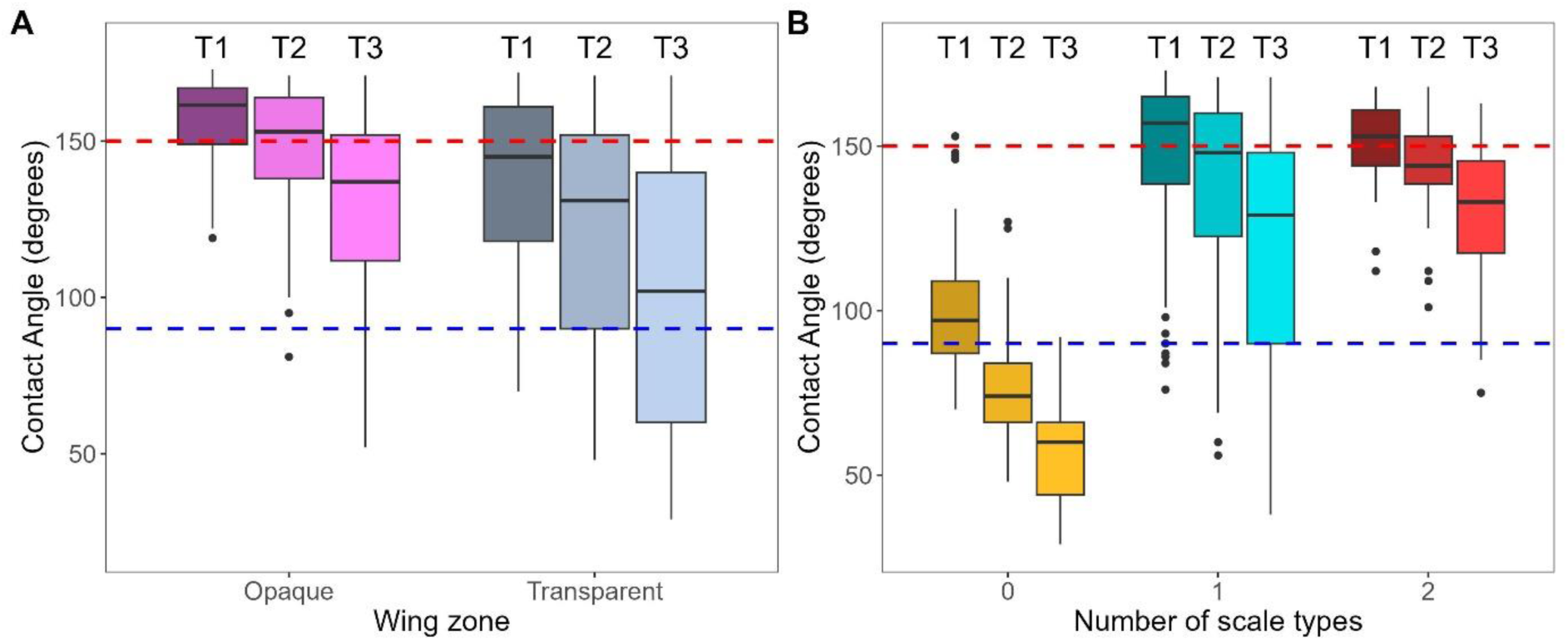
(A) Variation in contact angle with wing zone and time. (B) Variation in contact angle (A) with evaporation time and number of different scale types (0=nude membrane, 1=piliform or lamellar scales, 2= piliform and lamellar scales). All measurements and both zones were included. Time corresponds to water droplet size (T1: droplet of 1µl, T2: diameter divided by 2 relative to T1, T3 diameter divided by 4 relative to T1). Superhydrophobic: >150° (above the red line), hydrophobic: <150° and >90°; hydrophilic: <90° (below the blue line). Results are presented in ESM Tables S4a and S4b.

### Variation in hydrophobicity and relation to wing microstructure

The influence of wing microstructure on hydrophobicity is pervasive in our results (Figure 3). First, we find a higher interspecific variance in contact angle values in the transparent than in the opaque zone (Figure 2A, Fligner-Killeen tests with all times together χ²=79.48, p<0.001 or separated at T1: χ²=49.57, p<0.001; T2: χ²=29.24, p<0.001; T3 χ²=26.47, p<0.001), likely in relation to the higher interspecific microstructural diversity of the transparent zone. Second, the nude membrane (N) yields a lower hydrophobicity (positive scale presence effect in Figure 2B and ESM Table S4a) and a higher loss of hydrophobicity with water evaporation compared to the structural strategies that involved scales (Figure 2B and 3, positive scale presence x time interaction effect in ESM Table S4a). Third, presenting two types of scales (piliform and lamellar) or only one (piliform or lamellar) yields comparable levels of hydrophobicity (non-significant Scale Nb effect in ESM Table S4b). Yet, combining two types of scales attenuates the loss of hydrophobicity with water evaporation and improves the self-cleaning ability more than having only one type of scales only (Figure 2B and Figure 3, positive scale nb x time interaction effect in ESM Table S4b). Although erect and flat scales show comparable hydrophobicity (non-significant insertion effect in ESM Table S4b), erect scales limit more efficiently the loss of hydrophobicity with water evaporation than flat scales (Figure 3, negative insertion x time interaction effect in ESM Table S4b).

**Figure 3.**
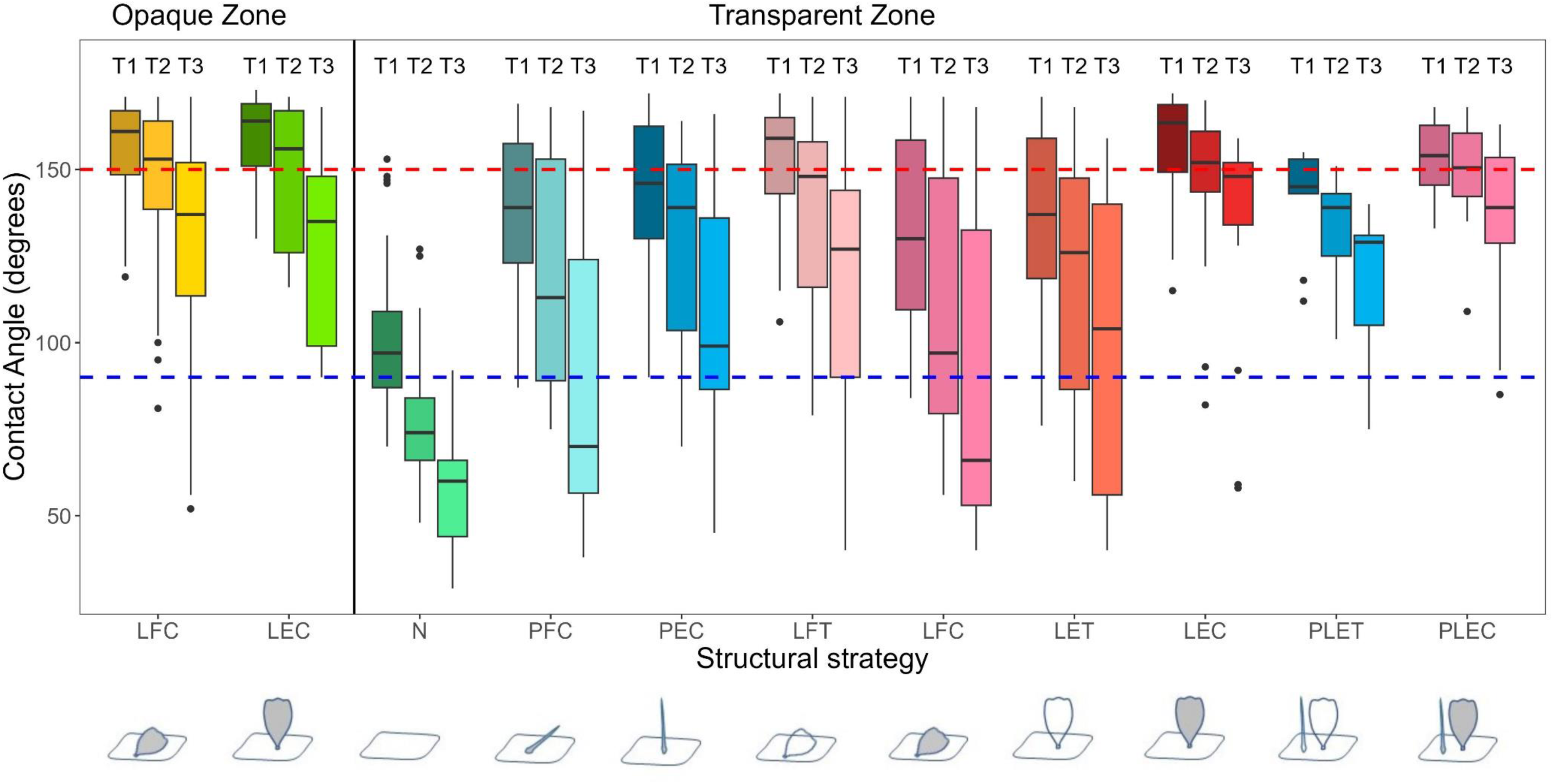
Variations of contact angle with wing zone, microstructure and time, i.e. water droplet size. Structural strategy is a combination of scale type (N: no scales, P: piliform scales, L: lamellar scales, PL: combination of piliform scales and lamellar scales), scale insertion (E: erected, and F: flat), and scale colour (C: coloured, and T: transparent). Superhydrophobic: >150° (above the red line), hydrophobic: <150° and >90°; hydrophilic: <90° (below the blue line). All individuals and droplets were considered. Results are presented in ESM Tables S2 to S3d.

Considering all scales together, transparent scales are less hydrophobic than coloured scales which are highly hydrophobic (negative scale colour (T>C) effect in ESM Table S4b). Yet, in detail, the effect of scale coloration depends on scale insertion. In flat scales, hydrophobicity is higher in transparent scales than in coloured scales (positive colour effect in ESM Table S4d), but it shows similar loss with water evaporation in transparent and in coloured scales (colour x time effect not retained in ESM Table S4c). In erect scales, hydrophobicity is similar in transparent and coloured scales (non-significant colour effect in ESM Table S4c) but transparent scales lose more hydrophobicity with water evaporation than coloured scales (negative colour x time effect in ESM Table S4c).

Considering all scales together, increasing scale density increases hydrophobicity (positive density effect in ESM Table S4b). Again, in detail, the effect of density depends on scale insertion. This effect is seen in flat scales (positive density effect in ESM Table S4d) but not in erect scales (non-significant density effect in ESM Table S4c). In erect scales, increasing scale density attenuates the loss of hydrophobicity with water evaporation (positive density x time interaction effect in ESM Table S4c, ESM Figure S3A) but it is not the case in flat scales (ESM Figure S3A). In flat scales, increasing the number of layers of scales attenuates the loss in hydrophobicity with water evaporation (positive NL x Time interaction effect in ESM Table S4d, ESM Figure S3B). Finally, the gain in hydrophobicity with increasing scale density is higher for transparent than for coloured scales (positive scale colour x density interaction effect in ESM Table S4b).

Given that erect geometries (involving piliform and/or lamellar scales: PLE, PE, LE) seem to interact differently with water compared to flat geometries, we analysed scale dimensions and spacing (ESM Table S5). In species with only lamellar or only piliform scales, we find that after controlling for wing size, erect piliform scales are thinner than flat piliform scales (ESM Figure S5C), and both have similar length (ESM Table S5CD, Figure S5A). Lamellar scales are shorter when erect than when flat, especially when they are colored rather than transparent (ESM Figure S5B, Table S5A). Lamellar scales show similar width whether they are flat or erect on the wing membrane and transparent lamellar scales are larger than colored lamellar scales (ESM Table S5B). In species with erect lamellar and piliform scales, piliform and lamellar scales are in similar densities (density ratio close to 1 for the intercept in ESM Table S5F, Figure S6B), and closely associated in space (insertion effect lower for E for spacing in ESM Table S5G, Figure S6C). In species with flat scales, piliform scales are not as dense as lamellar scales (insertion effect negative in ESM Table S5F) and more distantly associated in space (ESM Table S5G). These relationships are found while controlling for phylogeny, suggesting that the tight spatial association between piliform scales and lamellar scales in hierarchical PLE geometries is likely the result of selection. In species with both lamellar and piliform scales, piliform scales are 2.6 times longer than lamellar scales, creating a multi-hierarchical roughness at microscopic scales (ESM Figure S6A). In flat geometries, piliform scales are rare compared to lamellar scales, and both are more distantly spaced (ESM Table S5FG, Figure S6B); piliform scales are 5 times longer than lamellar scales (ESM Figure S6A).

### Variation in hydrophobicity in relation to optics

Using spectrometric measurements of wing direct transmittance, we find a negative relationship between contact angle and mean transmittance over 300-700 nm (Figure 4, ESM Table S6). A 10% increase in transmittance results in a 4° loss in CA. The relationship is statistically significant when considering all measurements or mean individual values, and marginally significant when considering mean species values, likely because of weaker statistical power.

**Figure 4.**
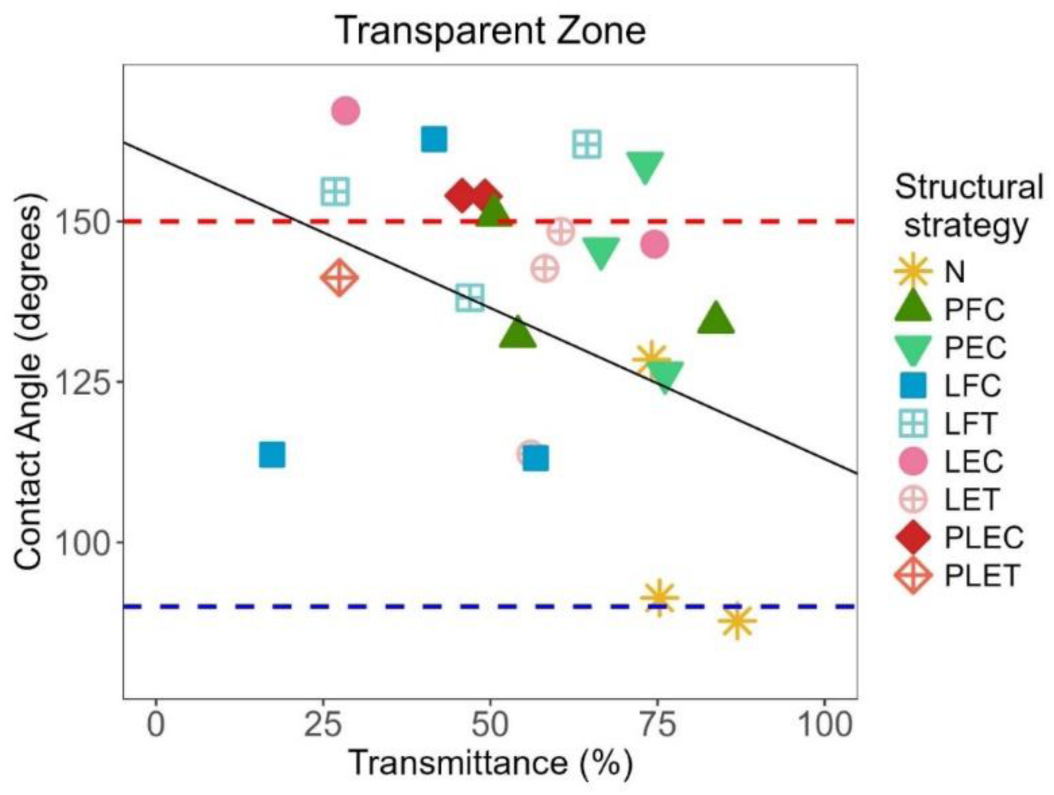
Variations of contact angle with wing transmittance for the different structural strategies. Structural strategy is a combination of scale type (N: no scales, P: piliform scales, L: lamellar scales, PL: combination of piliform scales and lamellar scales), scale insertion (E: erected, and F: flat), and scale colour (C: coloured, and T: transparent). Superhydrophobic: >150° (above the red line), hydrophobic: <150° and >90°; hydrophilic: <90° (below the blue line). We considered only the mean of CA for each species, for time T1, and for the transparent zone. The black plain line indicates the significant fitted regression line based on the Bayesian model. Results are presented in ESM Table S7.

### Variation in hydrophobicity in relation to the environment

Finally, compared to their temperate counterparts, species living in the tropics have a higher hydrophobicity in their transparent zone – loss of 10° CA for 10° increase in latitude – but a similar hydrophobicity in their opaque zone (Figure 5, ESM Table S7). All species show superhydrophobic opaque patches (Figure 5B, intercept above 150° in ESM Table S7B). There is no relationship between the proportion of wing area occupied by transparency and latitude that can have explained the observed variations in CA (ESM Table S7C).

**Figure 5.**
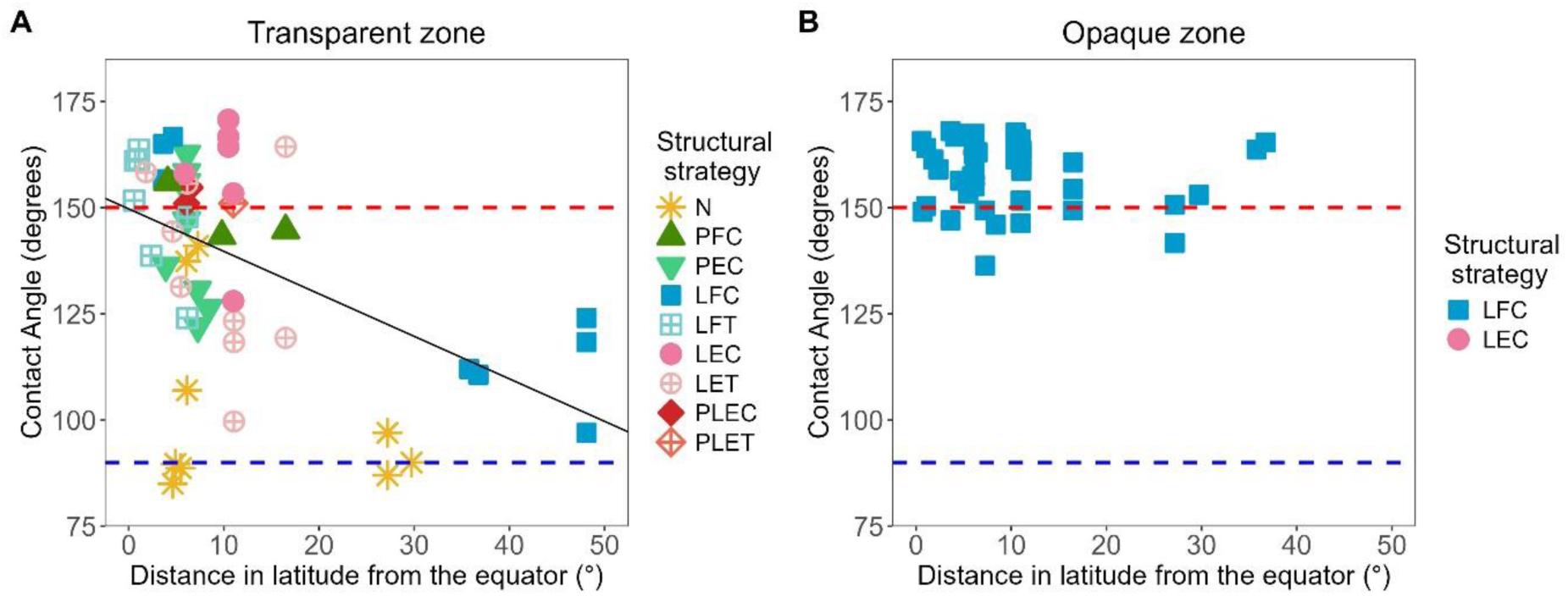
Relationship between contact angle in the transparent (A) and in the opaque (B) zone and the distance in latitude to the equator. Structural strategy is a combination of scale type (N: no scales, P: piliform scales, L: lamellar scales, PL: combination of piliform scales and lamellar scales), scale insertion (E: erected, and F: flat), and scale colour (C: coloured, and T: transparent). Superhydrophobic: >150° (above the red line), hydrophobic: <150° and >90°; hydrophilic: <90° (below the blue line). The black plain line in A indicates the significant fitted regression line based on the Bayesian model. Results are presented in ESM Table S8.

## DISCUSSION

### Variation in hydrophobicity and relation to wing macrostructure

We provide evidence for the first time at a broad taxonomic level that transparency is costly in terms of water repellency in Lepidoptera: transparent patches are less hydrophobic than opaque patches. We show that transparent patches have a lower hydrophobicity than opaque patches, and lose more hydrophobicity with water evaporation than opaque patches. Transparency thus entails important costs in terms of hydrophobicity in these two aspects. A loss of hydrophobicity with water evaporation has been commonly observed in hydrophobic human-made surfaces (McHale et al., 2005; Reyssat et al., 2007; P. Tsai et al., 2010) and in natural surfaces, as in the transparent-winged damselfly *Ischnura heterosticta* (Hasan et al., 2012). It has been interpreted as a loss of self-cleaning ability, especially when contact angle values get below the hydrophilicity threshold (Hasan et al., 2012).

We do not find any correlation between hydrophobicity and wing length, wing area to volume ratio, or forewing elongation, which at first sight partly contrasts with previous findings. In broad analyses covering many insect orders, Wagner et al. (1996) have found a positive correlation between CA and the ratio of wing area to body mass while Byun et al. (2009) have found a marginal positive correlation between CA and wing length and no correlation between CA and LW ratio (see ESM for analyses of their dataset). Yet, restricting their datasets to species more similar to Lepidoptera (with both wings involved in flight for Wagner, for wings with similar LW ratios in Byun), these relationships disappear (see ESM). More interestingly, we find that species with smaller wing area relative to their body or with shorter wings exhibit a greater loss of hydrophobicity with evaporation, hence lower self-cleaning ability. Species with a large body mass or short wings may move wings faster and small water droplets may roll off easily just through movement, attenuating selection for a high hydrophobicity towards small water droplets. The fact that species with more elongated wings exhibit a greater loss of hydrophobicity with evaporation (hence a lower self-cleaning ability) is more in contradiction with these results. A study of behaviour is needed to investigate further the relationships between wing macrostructure and water repellency ability.

### Variation in hydrophobicity and relation to wing microstructure

Our results show close relationships between transparency and microstructure and demonstrate that variations in microstructure can mitigate the costs of transparency.

Going back to the physical theory behind hydrophobicity, several studies have shown that a single-level structure does not necessarily guarantee a low water adhesion, even in the Cassie-Baxter state (see references in Su et al., 2010). Introducing higher levels of hierarchy increases the robustness of a surface hydrophobicity (Bell et al., 2015): it stabilizes the Cassie-Baxter state by dramatically decreasing the contact area fraction (ratio of contact area to the total surface area of the structure) and thus the adhesion force of water droplets, and by enlarging the energy barrier between the Cassie-Baxter and the Wenzel states. Hierarchical structures can be frequently found in plants and in animals. For instance, in the water strider *Gerris remigis*, leg water resistance is due to the hierarchical structures of nano-grooved microsetae, which prevented striders from being drowned under heavy rainfall (Gao & Jiang, 2004). A similar combination of micro- and nano-structuration has been found in the legs of mosquitoes, which ensured high hydrophobicity and high water-supporting ability; as a result, mosquitoes could stand effortlessly and walk easily on water (Wu et al., 2007). This likely explains why, in our study, the combination of erect piliform scales and lamellar scales yields a better self-cleaning property than piliform or lamellar scales alone. Interestingly, such geometries have a 3-level roughness: (1) erect piliform scales bending over lamellar scales (piliform scales are 2.6 times longer than lamellar scales and first in contact with water), (2) erect lamellar scales tightly associated in space with piliform scales (similar density and close spacing), and (3) nanostructures on scales and on the wing membrane. Hydrophobicity and self-cleaning likely results from the combination of the complex geometry of erect microstructures (which considerably reduces the proportion of the total surface in contact with water), and the gain in mechanical resistance (gain in elasticity and resistance against breakage) of piliform scales when bending against lamellar scales.

The importance of elasticity of bending hair-like microstructures has been found in several cases. In *Malacosoma castrensis* moths living by the sea, caterpillars withstood several hours to being flooded through a plastron protected by hairs (Kovalev et al., 2020). In the Lady’s mantle plant (*Alchemilla vulgaris*), hairs were hydrophilic when measured individually, but they bended and coalesced into bundles when in contact with water droplets; their elasticity resulted in a repulsive interaction between the droplet and the plant surface, which maintained hydrophobicity (CA above 90°) (Otten & Herminghaus, 2004). Likewise, in *Nasutitermes* termits, large bending hairs and small micrasters (micraster wavelength was around 11,7 µm according to our measurements taken on Figure 4C from 41) enabled hydrophobicity (CA above 90°) in both rain and mist conditions (Watson et al., 2011). Finally, in mosquitoes, the huge buoyant force developed by the legs ensuring easy movement on water largely stemmed from their mechanical flexibility (Kong et al., 2015). The extent of scale elasticity in Lepidoptera and its contribution to hydrophobicity needs specific experimental study.

Increasing scale density helps water repellency, regardless of the type of scales. In erect scales, increasing scale density does not influence CA but it attenuates the loss of hydrophobicity, thus improving self-cleaning ability. This latter result contradicts a previous finding that increasing the density of erect pillars increased the loss of hydrophobicity (Reyssat & Quéré, 2009). However, it is likely that increasing density stabilises the structure and increases the energy barrier between Cassie-Baxter and Wenzel regimes, resulting in maintaining high hydrophobicity despite water evaporation. In flat scales, increasing scale density increases hydrophobicity. In addition, increasing scale overlapping (number of layers) attenuates the loss of hydrophobicity with evaporation and improves self-cleaning. In the literature, scale overlap was assumed to help anisotropy in hydrophobicity (Bixler & Bhushan, 2014). In other words, during droplet evaporation, the Cassie-Baxter regime is more robust for denser microstructures. The mechanism by which hydrophobicity is maintained even for small droplets in multiple scale layers is still puzzling, but it may be related to scale arrangement, more specifically to scale bending, or to scale fine ridge ultrastructure (Burdin et al., 2025). Further experimental and modelling research is needed to clarify this density effect.

Not only scale architecture but also coloration can contribute to hydrophobicity. Erect scales show a lower loss of hydrophobicity (hence a greater self-cleaning ability) when pigmented than when transparent. In the transparent zone, coloured scales exhibit colours ranging from pale yellow to brown and black. They are likely impregnated by melanin pigments, which are known to be involved – for some biochemical forms – in cuticle sclerotization (hardening) (Sugumaran, 2009). Hence, the additional hardening conferred by pigments may increase their mechanical resistance to deformation and may contribute to maintaining hydrophobicity, even when evaporation occurs. The fact that in the flat geometry coloured and transparent scales lead to similar properties indicates that the role of colouration on hydrophobicity is more likely related to a change of elastic properties of the scales than a change of their surface chemistry.

Wing mechanical resistance is crucial for flight and geometries that limit protrusion height are more resistant to breakage while maintaining hydrophobicity (Bittoun & Marmur, 2012). Several of our results suggest scale height may be limited: (i) when piliform scales are alone, they have similar height, be they flat or erect, likely because they bend easily, which may limit their sensitivity to breakage. (ii) Erect lamellar scales are shortened and widened compared to flat lamellar scales, which likely increases their resistance to breakage. (iii) Erect transparent lamellar scales are densely packed, as shown in Gomez et al. (2021), which can also increase their mechanical resistance.

Our results bring novel evidence for a major role of microstructures in explaining large variations in hydrophobicity when diverse microstructures are considered. The rare existing studies on the subject suggest a synergetic effect of scale nanostructures and microstructures on enhancing surface hydrophobicity (experiments on one type of microstructure, namely flat lamellar scales in opaque butterflies (Fang et al., 2015; Aideo & Mohanta, 2021) or hairs in the wing of the housefly *Musca domestica* (Wan et al., 2019); theoretical modelling on one type of microstructure (Sajadinia & Sharif, 2010)), or even a major role of nanostructures in the overall variation (Fang et al., 2015; Wan et al., 2019). Yet, these two latter analyses only examined one type of microstructure, thereby potentially underestimating their contribution relative to that of nanostructures that were the only parameters that varied between their study species. Further experiments are needed to elucidate these aspects, and clarify the role of nanostructures, as not only their presence, but their topography and its randomness have been recently suggested to play a role in determining antiwetting properties (Li et al., 2020). Our study also shows that the elastic properties of the microstructures plays a significant role.

### Trade-off between hydrophobicity and optical transparency

In agreement with our prediction that microstructures play a major role in hydrophobicity, we find a negative relationship between hydrophobicity and transparency, a condition associated with major modifications in scale shape and density. This trade-off can be seen from the literature: the nymphalid butterfly *Greta oto* has been shown to exhibit a high transparency resulting from a low density of erect piliform scales and highly antireflective nanostructures (Pomerantz et al., 2021; Siddique et al., 2015) but a weak hydrophobicity (Wanasekara & Chalivendra, 2011). Likewise, the trade-off can be seen in the dragonfly *Gynacantha dravida* (which has micro and nanospikes), in which distal wing parts show higher hydrophobicity but lower transmittance compared to proximal wing parts (Aideo & Mohanta, 2016).

Despite the trade-off between optical transparency and water repellency, the nude membrane, which shows the highest optical transparency and the lowest hydrophobicity, maintains a weak hydrophobicity or is hydrophilic. In such species, where wings are deprived of scales, membrane nanostructures are at full play. Membrane nanostructures reduce reflection, but their efficiency at reducing water adhesion depends on the species. The nipple array maintains a highly hydrophobic surface in the cicada *Aleeta curvicosta* (CA=144° in Watson et al., 2008) but it fails at maintaining hydrophobicity in the hesperiid *Phanus vitreus* once erect transparent scales are removed (CA=92.8° in Finet et al., 2023). The variable efficiency of nanostructures at repelling water may depend on their other parameters - density, shape, spatial disorder – calling for detailed study of those features. Questions are still open regarding the role of randomness in nanostructures, shown to improve optical transparency (Siddique et al., 2015) but suggested to impair hydrophobicity (M. Sun et al., 2012; but see Li et al., 2020).

Revealing a trade-off between different properties and functions shows that species are submitted to antagonistic needs but can mitigate the ecological costs of clear wings. Several microstructural strategies – involving piliform and or lamellar scales, flat or erect, coloured or transparent – can show similar optical properties and levels of light transmission through the wings (drawing a horizontal line in Figure 6 in Gomez et al., 2021). Yet, these similarly optically-efficient microstructures differ in their hydrophobic properties, the combination of piliform and lamellar scales being most efficient. Hence, the high microstructural diversity (in scale presence, type, insertion, coloration, and density) allows species to offset some costs linked to transparency and tune functions separately, to a certain extent.

### Hydrophobicity and latitude

We find that tropical species have more hydrophobic transparent patches than temperate species, suggesting microstructural features are under selection. This result is consistent with the prediction that in tropical climates where species face more humid conditions, and where rainfall can happen daily, there is a stronger selective pressure for increased hydrophobicity. While the opaque zone allows maximizing hydrophobicity in all environmental conditions, the differential in environmental conditions reveals the costs of transparency. To our knowledge, this is the first evidence for a higher hydrophobicity in more humid conditions. Scarce relevant studies have explored the link between habitat humidity and species hydrophobicity: at local geographical scale, all four cicada species studied by (Oh et al., 2017) showed superhydrophobicity regardless of whether they live in dry or more humid habitats, but annual species were more hydrophobic than the species that emerges in large swarms every 17 years. Likewise, Goodwyn et al. (2009) suggested that in transparent butterflies hydrophobicity may depend on lifespan and migration ability.

Further studies are needed to elucidate the links between hydrophobicity and species ecology and disentangle the relative contributions of micro and nanostructures to wing hydrophobicity in Lepidoptera and exploring novel questions, like the role of randomness in structural organization.

## ACKNOWLEDGEMENTS AND FUNDING

We warmly thank Jacques Pierre, Rodolphe Rougerie, Thibaud Decaëns, Daniel Herbin, and Claude Tautel who helped with species choice and identification, and Edgar Attivissimo for additional Keyence imaging. This work was funded by Clearwing ANR project (ANR-16-CE02-0012), HFSP project on transparency (RGP0014/2016) and a France-Berkeley fund grant (FBF #2015-58).

## DATA AVAILABILITY STATEMENT

Data and R code are available on an OSF public repository at: https://doi.org/10.17605/OSF.IO/GB49E

## SUPPORTING INFORMATION

It includes one file called Gomez et al_SupportingInformation.docx

