## Supplementary text, tables and figures for "Stay clear and dry! How microstructure diversity can offset the hydrophobicity costs of transparency in clearwing Lepidoptera"

**ELECTRONIC SUPPLEMENTARY MATERIAL**

### METHODS

All code and data for these analyses are available with the article.

#### Hydrophobicity measurements

First, we performed a preliminary session of measurements to assess measurement repeatability. In a restricted sample of four species (list in ESM Table S1), we took 3 specimens per species, for which we measured the forewing and the hindwing. For each species and wing, we dropped three droplets in the transparent zone and three droplets in the opaque zone, on the ventral side of the wing. We measured the static contact angle between the water droplet and the wing surface, when the water droplet was dropped (time T1), when its height was approximately divided by two (time T2) and when it was divided by four (time T3) compared to its original height (ESM Figure S4). For each droplet, we took one photo at each time and analysed each photo twice independently, using the Keyence built-in software. Using the 'rptR' package (Stoffel et al., 2017), we built a model in which we included the species and zone as fixed factors, and the wing, the individual, the time, and the photo as random factors.

Second, we planned to perform our measurements on dry dead specimens, as it has been usually done in previous studies (Fang et al., 2015; Sun & Fang, 2015; Wagner et al., 1996; Wanasekara & Chalivendra, 2011). Yet, we were concerned that humidity could change the relative species behaviour regarding water repellency, i. e. that species may not rank in a similar way in dry and humid state. To test the potential effect of humidity on hydrophobicity, we selected a subset of five species that represented the most common microstructures. All species had lamellar flat coloured scales in their opaque zone (LFC). In their transparent zone, *Neocarnegia basirei* had no scales, *Rothschildia erycina* had piliform flat coloured scales (PFC), *Phanus vitreus* had lamellar erect colored scales (LEC), *Brevioleria oleria* had piliform erect scales, and *Methona curvifascia* had a combination of piliform and lamellar erect coloured scales (PLEC). We selected one specimen for each species and measured the contact angle in the opaque zone and in the transparent zone using the aforementioned protocol (3 water droplets per zone, at three times). We then moistened the specimens for 48h until specimens got fully rehydrated. We then repeated the measurements on the humid specimens. We then compared the series of CA values for dry and for humid treatment, and tested whether species ranking was conserved with humidity with a Spearman rank correlation in the opaque zone and in the transparent zone separately.

From all species, 21 were measured by JP, 7 were measured by CH. On five species (list in ESM Table S1), both observers independently analysed the same photos.

**Relationship between CA and ratio of wing area to body mass**

Wagner et al. (1996) studied the relationship between hydrophobicity and the ratio of wing area to body mass in insects. Their dataset was highly diverse, comprising 38 species from 14 insect orders. They found a positive correlation between CA and the ratio of wing area to body mass (estimate=11.09 $\pm$ 3.75, t=2.96, p=0.007). Yet, their dataset comprised many orders with no exposed wings, likely under different sets of selective pressures than insects with exposed wings. We thus re-analysed their data and restricted their dataset to insects with a microstructured membrane but no elytra (Odonata, Ephemeroptera, Lepidoptera, some Planipennia).

**Relationship between CA and wing length**

Byun et al (2009) studied the relationship between hydrophobicity and wing length (wrongly named wing area in their graph). Their dataset was diverse, comprising 24 species from 10 insect orders. They found a marginal correlation between CA and wing length (estimate=10.58$\pm$5.33, t=1.99, p=0.067) and no relationship between CA and LW ratio (estimate=5.41$\pm$7.86, t=0.68, p=0.503). The range of LW ratio was much higher (LW ratio: min=1.2, mean=3.6, max=9.3) than in our dataset (LW ratio: min=1.8, mean=2.2, max=3.2). We thus re-analysed their dataset and restricted the LW ratio range to our range. the effect of wing length disappears (estimate=2.42 $\pm$ 5.09, t=-0.47, p=0.652).

**RESULTS**

**Repeatability of CA measurements**

We found that the two contact angle measurements of the same photo gave highly repeatable values, that all droplets from the same wing and zone, and measured at the same time yielded highly repeatable values, that all individuals measured for the same species, zone and wing gave highly repeatable values, and that measurements from both wings in the same zone gave highly repeatable values (ESM Table S2).

**Effect of humidity on CA measurements**

The ranking between species for the hydrophobicity properties was conserved between the dry and the humid treatment, either in the opaque zone (spearman rho = 0.693, p=0.00549) or in the transparent zone (spearman rho=0.779, p=0.000988). In other words, whether specimens were dry or wet, the most hydrophobic species were the same, and the least hydrophobic species were the same. Based on these results, we established our protocol on dry specimens. For each species, we kept three individuals to maintain analysis power, but we measured only the forewing on the dorsal side. We dropped three droplets in the transparent zone and three droplets in the opaque zone (when it was large enough, which was not possible in 5/23 species, list in ESM Table S1); for each droplet, we took three photos (T1, T2, and T3) that we analysed once.

**Influence of the observer on CA measurements**

Considering these analyses as two repetitions, we found that contact angles were repeatable at the photo level (n=32, R=0.88±.137, p=0.0017). We then tested whether an observer could have a systematic bias in the measurement: in a linear mixed model with contact angle as the dependent variable, the droplet as a random factor and the observer as a potential fixed effect, we found that the best model was the null model, and that the observer did not explain a significant amount of contact angle variations (effect associated to p=0.17). Hence, we could pool all contact angle measurements without correcting for an observer influence.

**Analyses of datasets on macrostructural features**

Re-analysing Wagner et al's dataset, we restricted their dataset to insects with a microstructured membrane but no elytra (Odonata, Ephemeroptera, Lepidoptera, some Planipennia). The relationship between CA and the wing area to body mass index disappears (n=21 species (4 insect orders), estimate=0.15 $\pm$ 1.12, t=-0.14, p=0.895).

Re-analysing Wagner et al's dataset, we restricted their dataset to a range of LW ratio identical to ours. The relationship between CA and the wing length disappears (estimate=2.42 $\pm$ 5.09, t=-0.47, p=0.652).

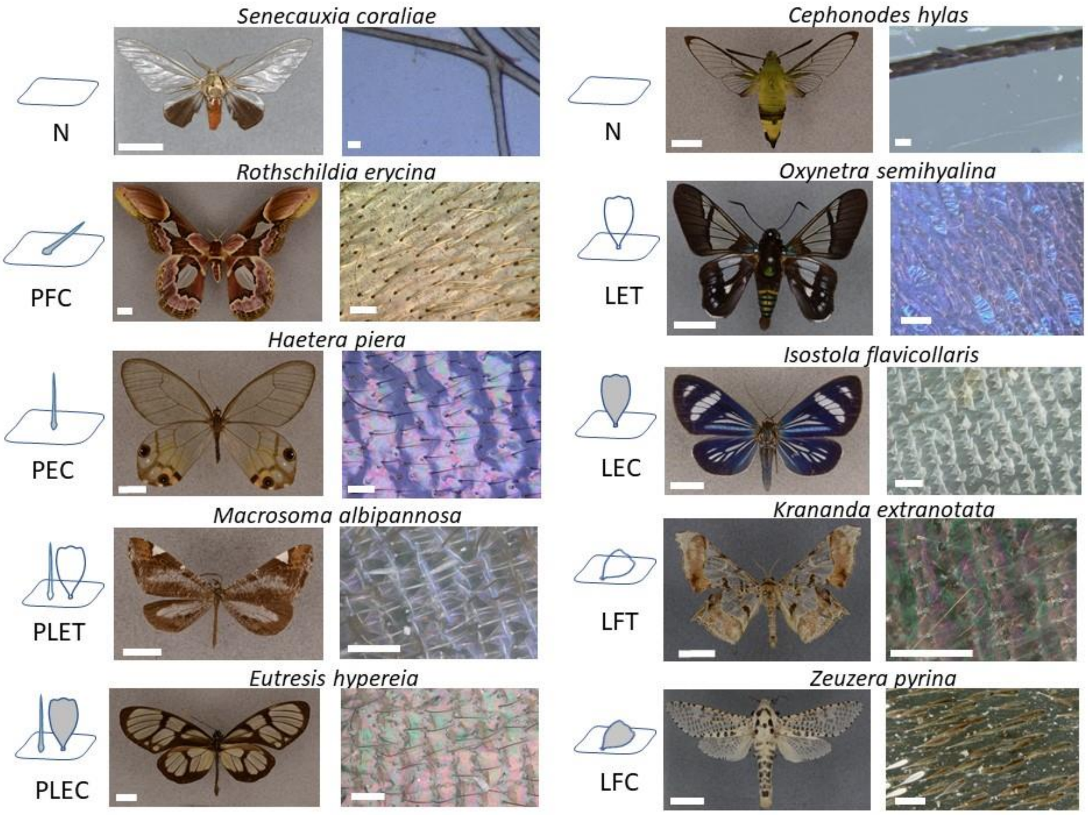


**Figure S1**. Examples of structural strategies in clearwing butterflies. Structural strategy is a combination of scale type (N: no scales, P: piliform scales, L: lamellar scales, PL: combination of piliform scales and lamellar scales), scale insertion (E: erected, and F: flat), and scale colour (C: coloured, and T: transparent). Notice that Macrosoma albipannosa has transparent lamellar scales but coloured piliform scales.

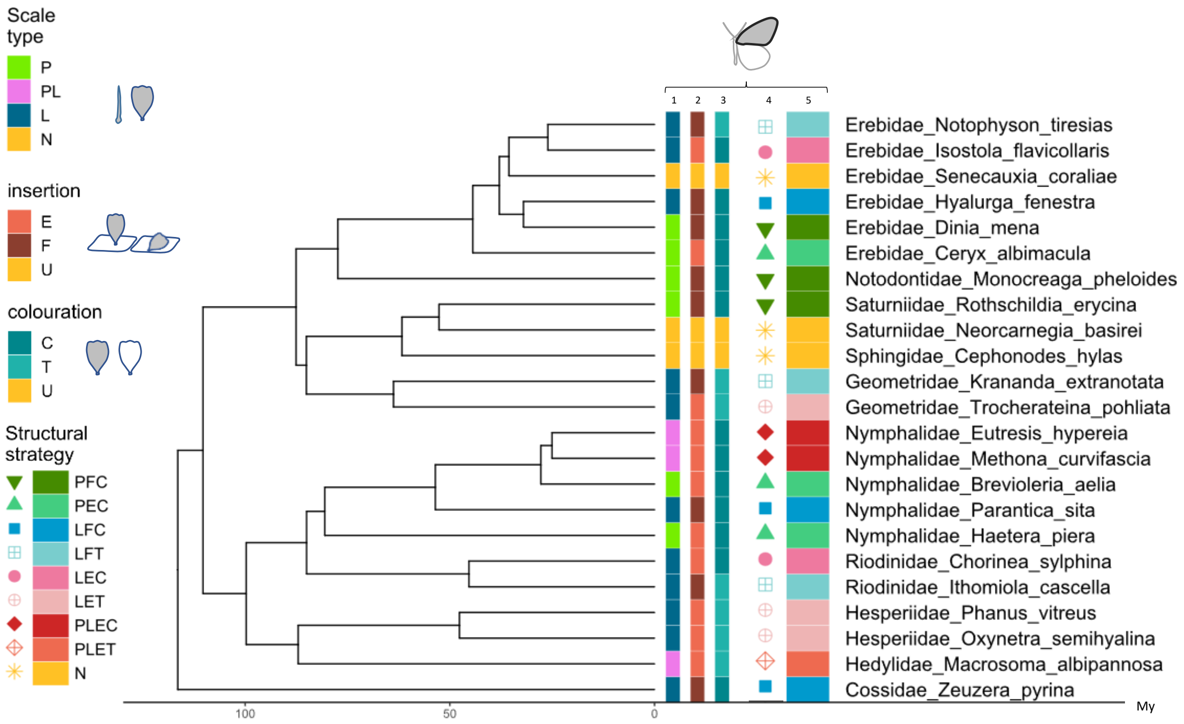


**Figure S2.** Phylogeny and distribution of trait values in the study species, for the forewing. Scale type (column 1), scale insertion on the membrane (column 2), scale colouration (column 3), and structural strategy (columns 4 and 5). For scale type: N=no scales, P=piliform scales, L= lamellar scales PL=combination of piliform scales and lamellar scales. For scale insertion on the membrane: E=erected, F=flat, U=undefined (for absent scales). For scale colouration: C=coloured, T=transparent, U=undefined (for absent scales). The strategy NUU was simplified into N.

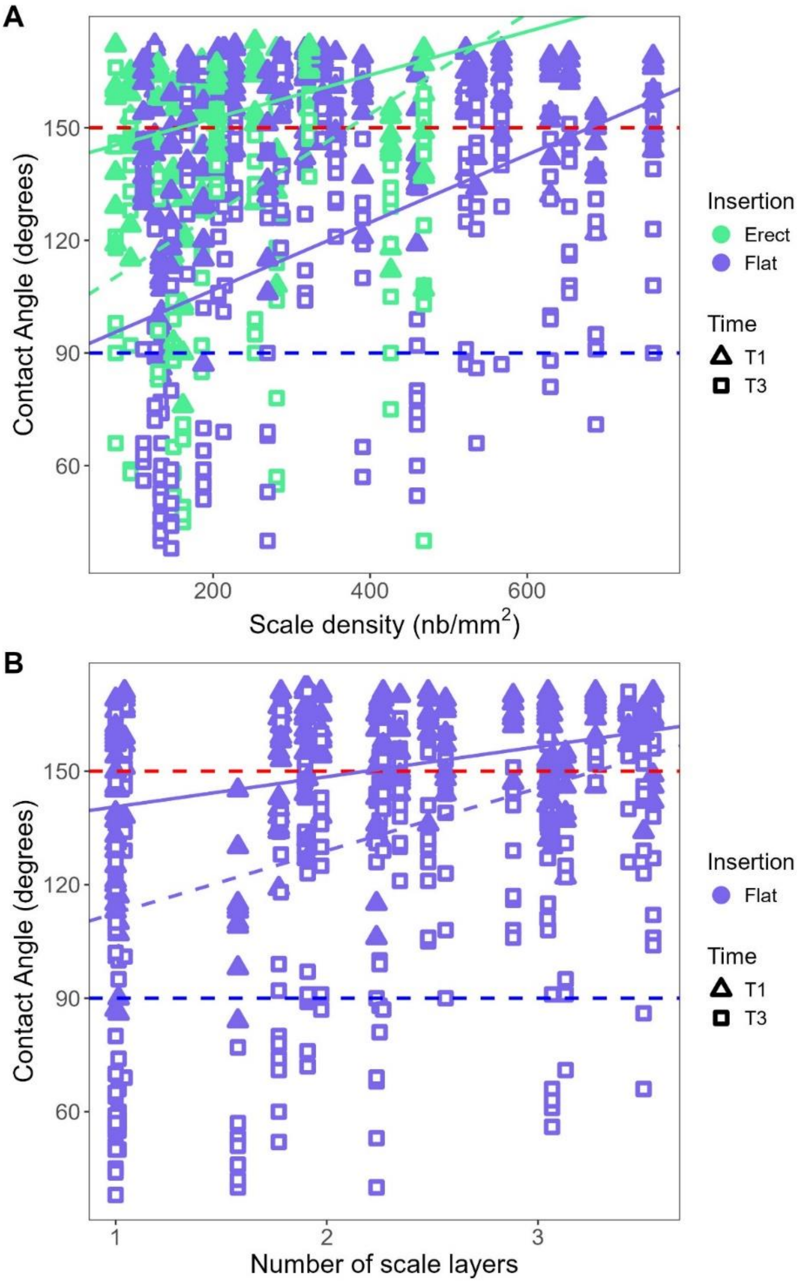


**Figure S3.** Variation in contact angle (A) with scale density for erect (green) and flat (violet) scales, (B) with the number of scale layers for flat scales. All measurements and both zones were included. Only T1 and T3 included. Superhydrophobic: >150° (above the red line), hydrophobic: <150° and >90°; hydrophilic: <90° (below the blue line). (B) regression lines are represented (solid line for T1, dashed line for T3 for erect scales, one line for flat scales as the interaction density x time was not significant. Results are presented in ESM Tables S4a, S4b for (A), S4c, S4d for (B), S4d for (C).

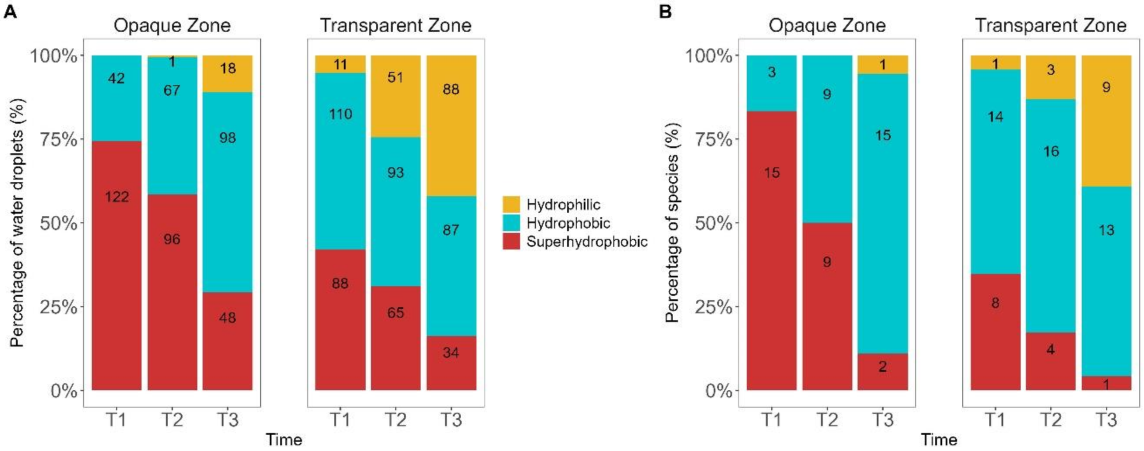


**Figure S4**. Percentage of water droplets (A) or species (B) that could be considered hydrophilic (CA<90°, gold), hydrophobic (90° ≤ CA ≤ 150°, blue), or superhydrophobic (CA>150°, brick), in the opaque zone and in the transparent zone, and at the different times T1, T2, and T3 measured. We took all the measured contact angle values for the water droplet level (A) while we took the mean contact angle values over all droplets and individuals for the species level (B). Numbers are presented per category within each stacked barplot.

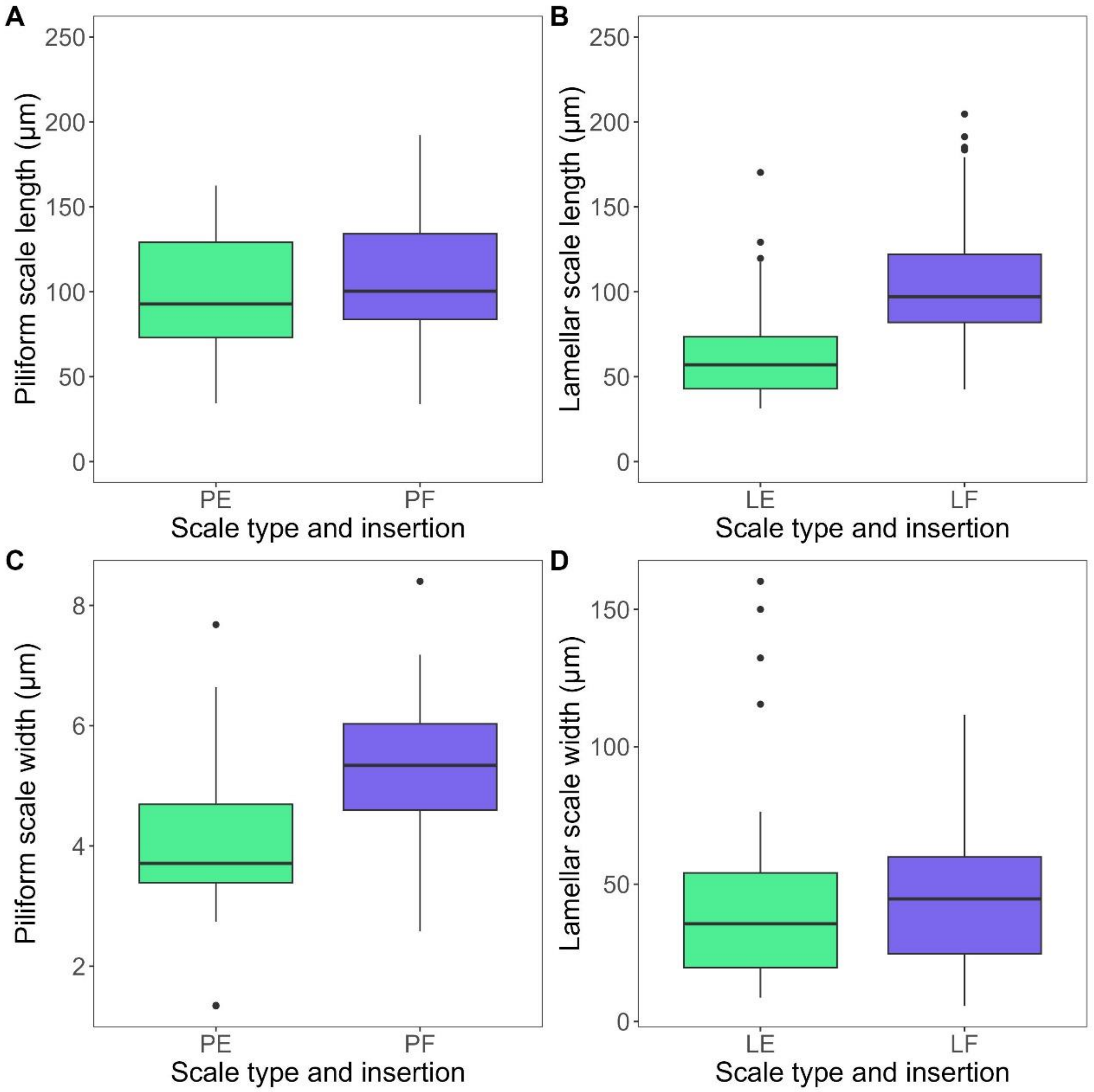


**Figure S5**. Geometries in structural strategies involving either piliform or lamellar scales alone, erect (purple) or flat (violet). Scale length for piliform (A) and lamellar scales (B), scale width for piliform scales (C) and lamellar scales (D) in relation to scale insertion, in the transparent zone. Symbols used are P=piliform scales, L=lamellar scales, E=erect, and F=flat. Analyses were done on 96 species of appropriate scale type and insertion and results are presented in ESM Table S6. 2 outliers were removed from (A) for clarity, 1 from (B).

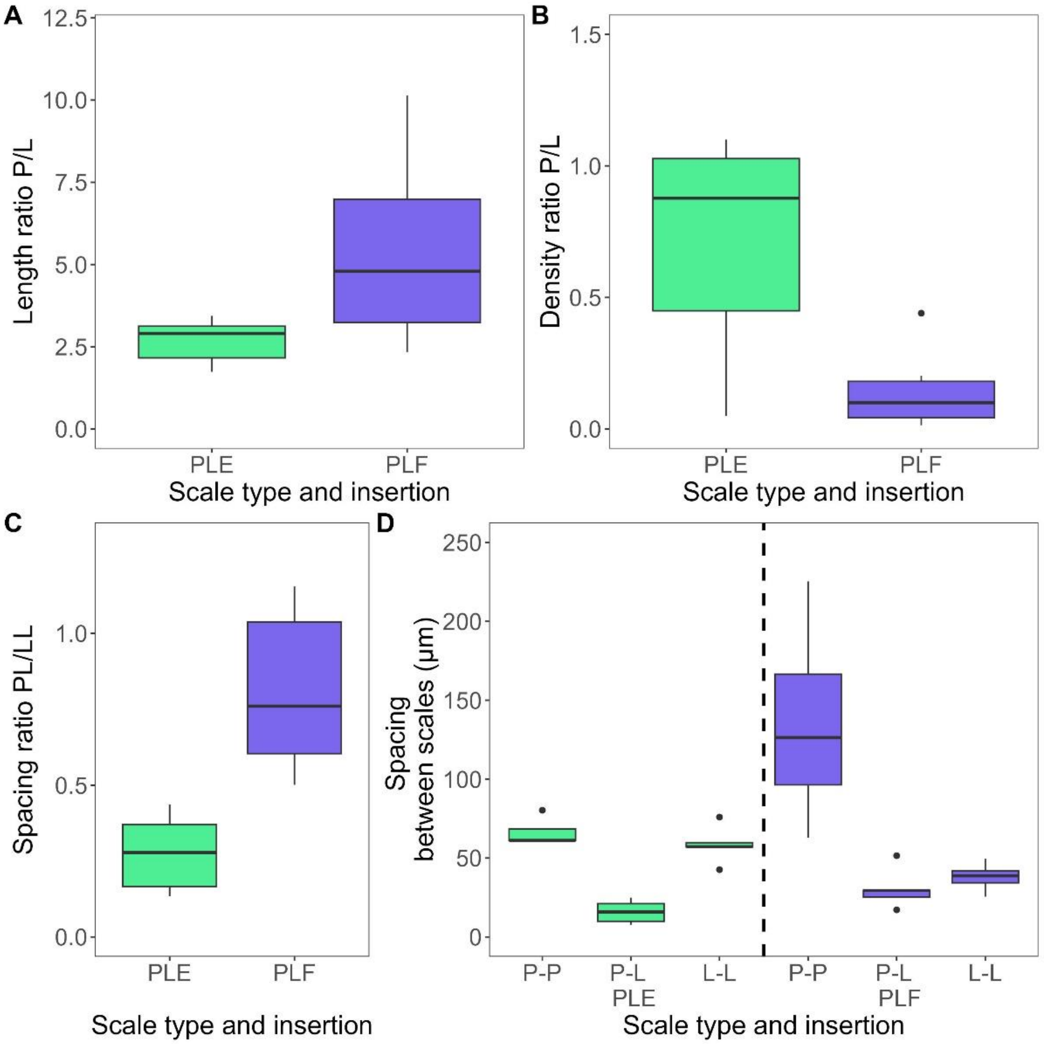


**Figure S6**. Geometries in structural strategies involving piliform and lamellar scales (PL strategies), erect (PLE, green) or flat (PLF, blue) in the transparent zone3,. Length ratio of piliform scale to lamellar scale (A), density ratio of piliform scale to lamellar scale (B), spacing ratio of PP/LL (C), and spacing between different types of scales (D) according to scale type and insertion, in the transparent zone. Symbols used are P=piliform scales, L=lamellar scales, E=erect, and F=flat. The 10 species analysed for PL strategy were *Athesis clearista, Diaphania unionalis, Dysschema boisduvalii, Eutresis hypereia, Macrosoma conifera, Macrosoma albipannosa, Methona curvifascia, Methona confusa, Nagara vitrea, Praeamastus fulvizonata*. Results are presented in ESM Table S6.

Table S1. List of species included in the study. Forewing structural strategy is a combination of scale type, insertion, and colouration. OZ=opaque zone, TZ=transparent zone. Scale type (N=no scales, P=piliform scales, L= lamellar scales, and PL=combination of piliform scales and lamellar scales), scale insertion (E=erect, F=flat), scale colouration (C=coloured, T=transparent). Species were measured by Jonathan Pairraire (JP) or Céline Houssin (CH). * *Macrosoma albipannosa* had transparent lamellar scales and coloured piliform scales. Q#1: is the protocol for measuring water repellency adapted? Q#2: does humidity affect species relative ranking for water repellency?

| SuperFamily | Family | SubFamily | Tribe | Species | Opaque Zone Measured | Tested for Q#1 | Tested For Q#2 | Observer | Forewing OZ structural strategy | Forewing TZ structural strategy |
|---|---|---|---|---|---|---|---|---|---|---|
| Bombycoidea | Saturniidae | Ceratocampinae | | Neorcarnegia basirei | yes | | yes | JP, CH | PFC | N |
| Bombycoidea | Saturniidae | Saturniinae | Attacini | Rothschildia erycina | yes | | yes | JP, CH | PFC | PFC |
| Bombycoidea | Sphingidae | Macroglossinae | Dilophonotini | Cephonodes hylas | no | | | JP | PFC | N |
| Cossoidea | Cossidae | Zeuzerinae | | Zeuzera pyrina | no | | | JP | PFC | LFC |
| Geometroidea | Geometridae | Ennominae | | Krananda extranotata | yes | | | JP | PFC | LFT |
| Geometroidea | Geometridae | Larentiinae | | Trocherateina pohliata | yes | | | JP | PFC | LET |
| Noctuoidea | Erebidae | Arctiinae | Syntomini | Ceryx albimacula | yes | | | JP | PFC | PEC |
| Noctuoidea | Erebidae | Arctiinae | Ctenuchini | Dinia mena | no | | | JP | PFC | PFC |
| Noctuoidea | Erebidae | Arctiinae | Pericopini | Hyalurga fenestra | yes | | | JP, CH | PFC | LFC |
| Noctuoidea | Erebidae | Arctiinae | Pericopini | Isostola flavicollaris | yes | | | JP | PFC | LEC |
| Noctuoidea | Erebidae | Arctiinae | Pericopini | Notophyson tiresias | yes | yes | | JP | PFC | LFT |
| Noctuoidea | Erebidae | Arctiinae | Phaegopterini | Senecauxia coraliae | no | | | JP | PFC | N |
| Noctuoidea | Notodontidae | Dioptinae | Dioptini | Monocreaga pheloides | yes | | | JP | PFC | PFC |
| Papilionoidea | Hedylidae | | | Macrosoma albipannosa | yes | | | CH | PFC | PLET* |
| Papilionoidea | Hesperiidae | | | Oxynetra semihyalina | yes | | | JP | PFC | LET |
| Papilionoidea | Hesperiidae | | | Phanus vitreus | yes | | yes | JP | PFC | LET |
| Papilionoidea | Nymphalidae | Danainae | Ithomiini | Brevioleria aelia | yes | yes | yes | JP | PEC | PEC |

| | | | | | | | | | | |
|---|---|---|---|---|---|---|---|---|---|---|
| Papilionoidea | Nymphalidae | Danainae | Ithomiini | Eutresis hypereia | yes | | | CH | PFC | PLEC |
| Papilionoidea | Nymphalidae | Danainae | Ithomiini | Methona curvifascia | yes | yes | yes | JP, CH | PFC | PLEC |
| Papilionoidea | Nymphalidae | Satyrinae | Haeterini | Haetera piera | no | yes | | JP | PFC | PEC |
| Papilionoidea | Nymphalidae | Danainae | Danaini | Parantica sita | yes | | | JP | PFC | LFC |
| Papilionoidea | Riodinidae | Riodininae | Riodinini | Chorinea sylphina | yes | | | JP, CH | PFC | LEC |
| Papilionoidea | Riodinidae | Riodininae | Riodinini | Ithomiola cascella | yes | | | JP | PFC | LFT |

Table S2. Repeatability of variables for macrostructure, contact angle, and colour measurements. For wing macrostructure, we tested whether individuals, measured once, were representative of their species. For nanostructure, we tested if height, spacing, and width (measured in multiple locations on scales in four species *Brevioleria aelia, Methona curvifascia, Notophyson Tiresias,* and *Haetera piera*) were representative of a species, zone, and scale type (piliform of lamellar). For contact angle, we tested if CA (measured twice for each photo independently) was representative of a species, wing (we measured both FW and HW), and zone (we measured both TZ and OZ, except for the forewing of *Haetera piera* where there was no opaque zone wide enough for measurement), then of individual code (I1, I2, I3), time (T1, T2, T3), and photo. For coloration, we tested whether the five spectra taken on one wing were representative of a species, and of a species and wing. p-values are significant for (* p<0.05; ** p<0.01; **** p<0.001).

| Condition | Variable | Species Nb/ Measurement Nb | Level | R (± se) |
|---|---|---|---|---|
| Question: are measurements repeatable at a certain level, in particular at species level. In other words, is an individual representative of its species? | | | | |
| Prediction: if measurement is repeatable, its R (repeatability value) is associated to a significant p-value. | | | | |
| Wing macrostructure | Width (mm) | 23/69 | Species | 0.974 (± 0.01)*** |
| | Length (mm) | 23/69 | Species | 0.982 (± 0.008)*** |
| | Forewing LWratio | 23/69 | Species | 0.923 (± 0.032)*** |
| | Forewing area | 23/69 | Species | 0.98 (± 0.006)*** |
| | Total wing area | 23/69 | Species | 0.981 (± 0.008)*** |
| | Ratio total wing area/body volume | 23/69 | Species | 0.894 (± 0.04)*** |
| Contact Angle | Contact Angle | 4/636 | SpeciesxWingxZone | 0.122 (± 0.053)*** |
| | Contact Angle | 4/636 | Individual | 0.052 (± 0.052)*** |
| | Contact Angle | 4/636 | Time | 0.139 (± 0.102)*** |
| | Contact Angle | 4/636 | Photo | 0.66 (± 0.101)*** |
| Colouration | Mean transmittance | 23/230 | Species | 0.863 (± 0.05)*** |
| Colouration | Mean transmittance | 23/230 | Species x Wing | 0.043 (± 0.024)*** |

Table S3. Variations of contact angle with wing macrostructure. The best random effect for the classic model was droplet nested within individual within species. Bayesian phylogenetically controlled mixed models for contact angle. TZ=transparent zone and OZ= opaque zone. The question investigated and the full model are written above the analysis. The best model was selected based on DIC minimisation and backward p-value selection. We tested three random factors: phylogeny, species, and specimen, and we retained the assemblage of random factors that minimized the DIC value. Fixed effect estimates with 95% credibility intervals excluding zero are indicated in bold, associated with a Bayesian P-value (*P < 0.05; **P < 0.01; ***P < 0.001).

| Associated Figure | Dependent variable | Parameter | Estimate [lower 95% CI, upper 95% CI] | Effective size | Bayesian P-value |
|---|---|---|---|---|---|
| | Question: does water repellency ability and its change with water evaporation depend on wing macrostructural features?<br>Full model : CA~LWratio + TotalWingArea + WingAreaBodyVolume + WingLength + Zone + TimeNum + LWratio x TimeNum + TotalWingArea x TimeNum + WingAreaBodyVolume x TimeNum + WingLength x TimeNum + Zone x TimeNum<br>3 random factors tested: Phylogeny, Species, Specimen | | | | |
| | | Intercept | 167.63 [144.46, 192.10] | 3960 | 0.0003*** |
| | | Forewing LWratio | -3.52 [-11.19, 3.68] | 4306 | 0.374 |
| | | WingArea/BodyVolume | -3.12 [-10.90, 5.03] | 3960 | 0.446 |
| | | Wing Length | 0.38 [-9.34, 9.29] | 3960 | 0.934 |
| Fig 2<br>Fig 3 | | **Zone (TZ>OZ)** | **-12.03 [-18.87, -5.38]** | **3960** | **0.001**** |
| | | **Time** | **-14.50 [-16.92, -12.21]** | **3960** | **0.0003***** |
| | CA | **Forewing LWratio x Time** | **-2.65 [-4.23, -1.04]** | **3960** | **0.001**** |
| | | **WingArea/BodyVolume x Time** | **1.42 [-0.17, 2.94]** | **3960** | **0.075~** |
| | | **Wing Length x Time** | **1.69 [0.14, 3.27]** | **3910** | **0.038*** |
| | | **Zone (TZ>OZ) x Time** | **-3.02 [-6.07, 0.07]** | **3960** | **0.056~** |
| | | Phylogenetic variance | 755.80 [268.32, 1369.87] | 3960 | - |
| | | Specimen | 109.80 [52.96, 168.42] | 3917 | - |
| | | Residual variance | 462.03 [422.82, 502.42] | 4609 | - |

Table S4a. Bayesian mixed models for the effect of scale presence, in the transparent zone (TZ) as a function of scale presence (N for nude membrane, Y for all other cases). The best model was selected based on minimisation of DIC value and backward p-value selection. The three random factors : phylogeny, Species, Specimen were tested and the models with 3 random factors and 2 (phylogeny, Specimen) performed equally. Bold values are statistically important factors associated with 95%CI excluding zero in Bayesian models (associated to p-values (* p<0.05; ** p<0.01; **** p<0.001). The question investigated and the full model are written above the analysis.

| Asso-ciated Figure | Dependent variable | Parameter | Estimate [lower 95% CI, upper 95% CI] | Effective size | Bayesian P-value |
|---|---|---|---|---|---|
| | Question: does water repellency ability and its change with water evaporation depend on scale presence?<br>Full model : CA~ Scale presence (N<Y)+Time + Scale presence (N<Y) x Time<br>3 random factors tested: Phylogeny, Species, Specimen | | | | |
| Fig 3<br>Fig S3A | CA in species with or without scales | Intercept | 115.25 [76.14, 153.60] | 3500 | 0.0003*** |
| | | **Scale presence (N<Y)** | **39.78 [8.95, 71.64]** | **3500** | **0.011*** |
| | | **Time** | **-22.88 [-28.06, -17.85]** | **3500** | **0.0003***** |
| | | **Scale presence (N<Y) x Time** | **5.67 [-0.01, 10.97]** | **3462** | **0.043*** |
| | | Phylogenetic variance | 827.91 [268.05, 1550.84] | 3500 | - |
| | | Specimen | 223.49 [125.61, 352.70] | 3676 | - |
| | | Residual variance | 374.53 [334.40, 422.10] | 3500 | - |

Table S4b. Bayesian mixed models for the effect of the number of scale types (1 for piliform or lamellar scales, 2 for both piliform and lamellar scales combined). The question investigated and the full model are written above the analysis. We present the best model, selected through DIC minimization and backward p-value selection. We tested three random factors : phylogeny, species identity, and specimen identity and we retained the assemblage of random factors that minimized DIC value or, if with similar DIC value, the simpler assemblage. Bold values are statistically important factors associated with 95%CI excluding zero in Bayesian models associated to p-values (* p<0.05; ** p<0.01; **** p<0.001); less important factors are associated with 90%CI excluding zero (with symbol ~). The question investigated and the full model tested are written above the analyses.

| Associated Figure | Dependent variable | Parameter | Estimate [lower 95% CI, upper 95% CI] | Effective size | Bayesian P-value |
|---|---|---|---|---|---|
| | Question: does water repellency ability and its change with water evaporation depend on scale microstructural characteristics (number of different types, density, number of layers, insertion, and coloration?<br>Full model : CA~ ScaleNbType + ScaleDensity + ScaleInsertion + ScaleColour + NL + Time + WingLength + ScaleNbType x Time + ScaleNbType x ScaleDensity + ScaleDensity x Time + ScaleColour x Time + ScaleColour x Scale Density + NL x Time + ScaleNbType x Time x ScaleDensity + ScaleInsertion x Time x ScaleDensity + ScaleColour x Time x ScaleDensity<br>3 random factors tested: Phylogeny, Species, Specimen | | | | |
| Fig 2A<br>Fig 3 | CA<br>in all scales | Intercept | 157.92 [139.96, 176.80] | 4500 | 0.0002*** |
| | | Scale Nb (2>1) | -1.76 [-16.01, 13.54] | 5044 | 0.809 |
| | | **Time** | **-13.43 [-16.80, -9.98]** | **4218** | **0.0002*** ** |
| | | NL | 2.82 [-1.82, 8.03] | 4500 | 0.273 |
| | | Scale Insertion (F>E) | 1.46 [-9.09, 12.74] | 4274 | 0.797 |
| | | **Scale Colour (T>C)** | **-5.63 [-10.61, -0.72]** | **4286** | **0.027*** |
| | | **Scale Density** | **4.86 [1.58, 8.32]** | **3862** | **0.004**** |
| | | **Scale Nb (2>1) x Time** | **6.47 [0.30, 13.13]** | **4500** | **0.047*** |
| | | **NL x Time** | **4.13 [1.91, 6.25]** | **4500** | **0.0002*** ** |
| | | **Scale Insertion (F>E) x Time** | **-4.84 [-9.47, -0.60]** | **4070** | **0.034*** |
| | | **Scale Colour (T>C) x Scale Density** | **20.22 [12.88, 26.72]** | **4500** | **0.0002*** ** |
| | | Phylogenetic variance | 377.92 [110.27, 734.28] | 4500 | - |
| | | Specimen | 133.07 [70.16, 210.05] | 4500 | - |
| | | Residual variance | 428.84 [392.07, 468.51] | 4500 | - |

Table S4c. Effect of scale microstructural features for erect scales only. ET= erect transparent scales, and EC= erect coloured scales. The number of layers was 1 for all erect scales, hence we did not test that factor. Since they were almost only in the transparent zone, we could not incorporate Zone as a factor. In erect scales, there could be either 1 scale type or 2 scale types, so we could test the effect of ScaleNbType. We present the best model, selected through DIC minimization and backward p-value selection. We tested three random factors : phylogeny, species identity, and specimen identity and we retained the assemblage of random factors that minimized DIC value or, if with similar DIC value, the simpler assemblage. Bold values are statistically important factors associated with 95%CI excluding zero in Bayesian models associated to p-values (* p<0.05; ** p<0.01; **** p<0.001); less important factors are associated with 90%CI excluding zero (with symbol ~). The question investigated and the full model tested are written above the analyses.

| Associated Figure | Dependent variable | Parameter | Estimate [lower 95% CI, upper 95% CI] | Effective size | Bayesian P-value |
|---|---|---|---|---|---|
| | Question: does water repellency ability and its change with water evaporation depend on the number of scale types, scale density, and scale colour?<br>Full model : CA~ ScaleNbType + ScaleDensity + ScaleColour + Time + ScaleNbType*Time + ScaleDensity*Time + ScaleColour*Time<br>3 random factors tested: Phylogeny, Species, Specimen | | | | |
| Fig 3<br>Fig S3B | CA for erect scales | Intercept | 166.96 [149.58, 182.86] | 4046 | 0.0003*** |
| | | Scale density | 3.77 [-11.51, 19.47] | 3500 | 0.627 |
| | | **Time** | **-9.64 [-14.51, -5.42]** | **3500** | **0.0003*** ** |
| | | Colour ET>EC | -13.64 [-39.07, 10.24] | 3895 | 0.263 |
| | | **Scale density x Time** | **7.28 [2.06, 12.82]** | **3500** | **0.011*** |
| | | **Colour ET>EC x Time** | **-8.59 [-16.02, -1.13]** | **3328** | **0.030*** |
| | | Phylogenetic variance | 75.93 [0.0003, 345.44] | 3500 | - |
| | | Specimen | 256.15 [96.02, 442.84] | 3700 | - |
| | | Residual variance | 410.45 [340.96, 476.62] | 3500 | - |

Table S4d. Effect of scale microstructural features for flat scales only. FT= flat transparent scales, and FC= flat coloured scales. In flat scales, there was only one type of scales, so we did not test ScaleNbType. We present the best model, selected through DIC minimization and backward p-value selection. We tested three random factors : phylogeny, species identity, and specimen identity and we retained the assemblage of random factors that minimized DIC value or, if with similar DIC value, the simpler assemblage. Bold values are statistically important factors associated with 95%CI excluding zero in Bayesian models associated to p-values (* p<0.05; ** p<0.01; **** p<0.001). The question investigated and the full model tested are written above the analyses.

| Associated Figure | Dependent variable | Parameter | Estimate [lower 95% CI, upper 95% CI] | Effective size | Bayesian P-value |
|---|---|---|---|---|---|
| | Question: does water repellency ability and its change with water evaporation depend on scale characteristics for flat scales in particular?<br>Full model : CA~ ScaleColour + ScaleDensity + NL + Time<br>+ ScaleDensity x Time + ScaleColour x Time+ ScaleDensity x ScaleColour+ NL x Time<br>+ ScaleDensity x Time x ScaleColour<br>3 random factors tested: Phylogeny, Species, Individual | | | | |
| Fig 3<br>Fig S3B<br>Fig S3C | CA for flat scales | Intercept | 151.00 [127.58, 175.04] | 3602 | 0.0003*** |
| | | **Scale Colour (FT>FC)** | **21.69 [9.67, 34.25]** | **4524** | **0.0003*****|
| | | **Scale Density** | **17.24 [11.42, 23.50]** | **4376** | **0.0003*****|
| | | NL | 3.65 [-0.95, 8.85] | 3900 | 0.138 |
| | | **Time** | **-18.35 [-20.25, -16.25]** | **3900** | **0.0003*****|
| | | **Scale Colour (FT>FC) x Scale Density** | **39.84 [20.60, 60.47]** | **3900** | **0.0003*****|
| | | **NL x Time** | **4.29 [2.17, 6.29]** | **3900** | **0.0003*****|
| | | Phylogenetic variance | 789.69 [240.52, 1567.87] | 3900 | - |
| | | Specimen | 120.65 [57.25, 197.68] | 3900 | - |
| | | Residual variance | 373.83 [333.72, 414.96] | 3900 | - |

Table S5. Variations of scale microstructural parameters in relation to scale geometry in the transparent zone. Bayesian phylogenetically controlled mixed models. Piliform scales (P), lamellar scales (L), piliform and lamellar scales (PL). Erect scales (E) and flat scales (F). Data came from the species from the large 123 species dataset analysed in Gomez et al. (2021) for scale length. For the other analyses, data came from 8 species: *Athesis clearista, Diaphania unionalis, Dysschema boisduvalii, Eutresis hypereia, Macrosoma conifera, Methona confusa, Nagara vitrea, Praeamastus fulvizonata.* We present the minimal models, selected through DIC minimization and backward p-value selection, keeping the same factors for models of scale length and scale width to facilitate comparison. We tested three random factors : phylogeny, species identity, and specimen identity and we retained the assemblage of random factors that minimized DIC value or, if with similar DIC value, that showed better convergence criteria. Bold values are statistically important factors associated with 95%CI excluding zero in Bayesian models associated to p-values (* p<0.05; ** p<0.01; **** p<0.001). The question investigated and the full model tested are written above the analyses.

| Asso-ciated Figure | Dependent variable | Sample size | Parameter | Estimate [lower 95% CI, upper 95% CI] | Effective size | Bayesian P-value |
|---|---|---|---|---|---|---|
| | A. Question: in lamellar scales, does scale length vary with scale insertion or coloration?<br>Full model : Scale Length~ Scale Insertion + ScaleColoration + WingLength + Scale Insertion x Scale Coloration<br>2 random factors tested: Phylogeny + Species | | | | | |
| Fig S5B | A. Scale length Lamellar scales | 69 spe | T | 52.76 [10.25, 100.02] | 4078 | 0.026* |
| | | | **Scale insertion (F>E)** | **48.58 [29.45, 68.88]** | **4121** | **0.0003*** ** |
| | | | Scale coloration (T>C) | 20.48 [-11.44, 52.46] | 3900 | 0.228 |
| | | | **Wing Length** | **1.05 [0.43, 1.74]** | **4157** | **0.003** ** |
| | | | **Scale insertion (F>E) x Scale coloration (T>C)** | **-39.59 [-73.27, -6.12]** | **3900** | **0.021*** |
| | | | Phylogenetic variance | 2060.26 [1161.39, 3132.01] | 4084 | - |
| | | | Residual variance | 412.17 [250.40, 594.67] | 4187 | - |
| | B. Question: in lamellar scales, does scale width vary with scale insertion or coloration?<br>Full model : Scale Width~ Scale Insertion* Scale Coloration+ Wing Length<br>2 random factors tested: Phylogeny + Species | | | | | |
| Fig S5D | B. Scale width Lamellar scales | 69 spe | Intercept | 30.90 [0.01, 60.00] | 4106 | 0.041* |
| | | | Scale insertion (F>E) | 5.12 [-5.59, 16.88] | 3900 | 0.376 |
| | | | **Scale coloration (T>C)** | **45.20 [26.31, 63.25]** | **3900** | **0.0003*** ** |
| | | | Wing Length | -0.08 [-0.43, 0.28] | 3900 | 0.622 |
| | | | Scale insertion (F>E) x Scale coloration (T>C) | -14.77 [-33.25, 4.77] | 3900 | 0.121 |

| | | | | | | |
|---|---|---|---|---|---|---|
| | | | Phylogenetic variance | 982.21 [590.22, 1395.50] | 3900 | - |
| | | | Residual variance | 96.56 [61.98, 141.05] | 3900 | - |
| | C. Question: in piliform scales, does scale length vary with scale insertion?<br>Scale Length~ Scale Insertion + Wing Length, because all piliform scales are coloured<br>2 random factors tested: Phylogeny + Species | | | | | |
| Fig S5A | C.<br>Scale length<br>Piliform scales | 27 spe | Intercept | 78.78 [30.42, 131.47] | 3500 | 0.006** |
| | | | Scale insertion (F>E) | 6.60 [-18.93, 35.91] | 3500 | 0.619 |
| | | | **Wing Length** | **1.55 [0.74, 2.34]** | **3500** | **0.001**** |
| | | | Phylogenetic variance | 2004.22 [0.0002, 4146.45] | 3500 | - |
| | | | Residual variance | 642.10 [273.61, 1066.39] | 1826 | - |
| | D. Question: does scale width vary with scale insertion?<br>Piliform scales: Scale width~ Scale Insertion + Wing Length, because all piliform scales are coloured<br>2 random factors tested: Phylogeny + Species | | | | | |
| Fig S5C | D.<br>Scale width<br>Piliform scales | 27 spe | Intercept | 3.80 [2.39, 5.47] | 3692 | 0.0003*** |
| | | | **Scale insertion (F>E)** | **1.37 [0.40, 2.26]** | **3500** | **0.002**** |
| | | | Wing Length | 0.02 [-0.01, 0.04] | 3386 | 0.223 |
| | | | Phylogenetic variance | 1.75 [0.0006, 3.84] | 3304 | - |
| | | | Residual variance | 1.00 [0.49, 1.63] | 3236 | - |
| | E. Question: does scale length ratio vary with scale insertion<br>Full model : Spacing~ Scale Insertion<br>2 random factors tested: Phylogeny + Species | | | | | |
| Fig S6A | E.<br>Length ratio<br>(=lengthP/ lengthL) | 8 spe | Intercept | 3.37 [-4.64, 10.27] | 4003 | 0.282 |
| | | | Scale insertion (F>E) | 2.58 [-4.68, 10.53] | 4274 | 0.411 |
| | | | Phylogenetic variance | 14.32 [0.0002, 40.18] | 3507 | - |
| | | | Species | 3.29 [0.0002, 14.17] | 2629 | - |
| | | | Residual variance | 0.18 [0.01, 0.54] | 4500 | - |
| Fig S6B | F. Question: does scale density ratio between piliform and lamellar scales vary with scale insertion<br>Full model : Scale Density Ratio~ Scale Insertion<br>2 random factors tested: Phylogeny + Species | | | | | |
| | F.<br>Density ratio | 8 spe | **Intercept** | **0.94 [0.56, 1.31]** | **3500** | **0.002**** |
| | | | **Scale insertion (F>E)** | **-0.80 [-1.17, -0.34]** | **3500** | **0.006**** |

| | | | | | | |
|---|---|---|---|---|---|---|
| | (= densityP/ densityL) | | Phylogenetic variance | 0.03 [0.0002, 0.11] | 3500 | - |
| | | | Residual variance | 0.05 [0.01, 0.10] | 3500 | - |
| | G. Question: does scale spacing vary with scale insertion and scale type?<br>Full model : Scale Spacing~ Scale Insertion + Scale Type<br>+ Scale Insertion x Scale Type ((PL= between piliform and lamellar scales, LL= between lamellar scales)?<br>2 random factors tested: Phylogeny + Species | | | | | |
| Fig S6C, Fig S6D | G. Scale spacing | 8 spe | Intercept | 37.89 [22.97, 52.45] | 7500 | 0.003** |
| | | | **Insertion (F>E)** | **21.68 [-0.84, 43.09]** | **7500** | **0.050~** |
| | | | Scale Type (PL>LL) | -7.43 [-21.76, 8.56] | 7500 | 0.307 |
| | | | **Insertion (F>E) x Scale Type (PL>LL)** | **-36.32 [-61.41, -10.31]** | **8239** | **0.009**** |
| | | | Phylogenetic variance | 60.99 [0.0002, 337.64] | 3730 | - |
| | | | Residual variance | 147.84 [31.30, 301.71] | 7171 | - |

Table S6. Variations of CA with the mean transmittance (over 300-700 nm) of the transparent zone. Bayesian phylogenetically controlled mixed models for contact angle in the transparent zone for T1 droplets, at droplet level, individual level, and species level. We present the best model, selected through DIC minimization and backward p-value selection. We tested three random factors : phylogeny, species identity, and specimen identity and we retained the assemblage of random factors that minimized DIC value or, if with similar DIC value, the simpler assemblage. Bold values are statistically important factors associated with 95%CI excluding zero in Bayesian models associated to p-values (* p<0.05; ** p<0.01; **** p<0.001); less important factors are associated with 90%CI excluding zero (with symbol ~). The question investigated and the full model tested are written above the analyses.

| Asso-ciated Figure | Dependent variable | Parameter | Estimate [lower 95% CI, upper 95% CI] | Effective size | Bayesian P-value |
|---|---|---|---|---|---|
| Fig 4 | | Question: Is there a trade-off between optical transparency and hydrophobicity, at droplet level, individual level, species level ?<br>Full model : Scale Length~ Scale Type + Scale Insertion + Scale Coloration + Wing Length + Scale Insertion x Scale Coloration+ Scale Insertion x Scale Type<br>3 random factors tested: Phylogeny, Species, Specimen | | | |
| | A. CA at droplet level | Intercept | 159.92 [125.57, 193.97] | 5758 | 0.0002*** |
| | | **[300-700] transmission** | **-0.46 [-0.89, -0.01]** | **5778** | **0.038*** |
| | | Phylogenetic variance | 847.75 [356.93, 1502.36] | 5500 | - |
| | | Specimen | 21.52 [0.0002, 77.07] | 4372 | - |
| | | Residual variance | 275.86 [210.94, 341.24] | 5257 | - |
| | B. CA at individual level | Intercept | 160.36 [125.41, 193.44] | 3806 | 0.0003*** |
| | | **[300-700] transmission** | **-0.47 [-0.90, -0.03]** | **3773** | **0.032*** |
| | | Phylogenetic variance | 828.11 [333.33, 1464.95] | 3900 | - |
| | | Residual variance | 136.01 [84.38, 198.47] | 3900 | - |
| | C. CA at species level | Intercept | 160.28 [128.49, 188.66] | 4500 | 0.0002*** |
| | | **[300-700] transmission** | **-0.41 [-0.88, 0.11]** | **4500** | **0.091~** |
| | | Phylogenetic variance | 80.59 [0.0002, 627.59] | 4032 | - |
| | | Species | 222.89 [0.0002, 665.83] | 5356 | - |
| | | Residual variance | 226.48 [0.0002, 646.73] | 3666 | - |

Table S7. Variations of CA with the distance in latitude to the equator. Bayesian phylogenetically controlled mixed models for contact angle, separately for the transparent zone (A) and for the opaque zone (B). The absolute distance in latitude from the equator was taken as the factor and included as the minimal model to test. We computed the models for T1 mean CA values for individuals for which we had GPS coordinates of their collection site. We present the best model, selected through DIC minimization and backward p-value selection. We tested two random factors : phylogeny, and species identity, and we retained the assemblage of random factors that minimized DIC value or, if with similar DIC value, the simpler assemblage. Bold values are statistically important factors associated with 95%CI excluding zero in Bayesian models associated to p-values (* p<0.05; ** p<0.01; **** p<0.001); less important factors are associated with 90%CI excluding zero (with symbol ~). The question investigated and the full model tested are written above the analyses.

| Associated Figure | Dependent variable | Parameter | Estimate [lower 95%CI, upper 95%CI] | Effective size | Bayesian P-value |
|---|---|---|---|---|---|
| | Question: Are species living in more humid areas (lower latitude) more hydrophobic than species in drier areas (higher latitude)? Full model : CA ~distance in latitude from the equator 2 random factors tested: Phylogeny, Species | | | | |
| Fig 5A | A. CA at individual level in the transparent zone | Intercept | 149.72 [136.59, 162.91] | 48616 | 0.00002*** |
| | | **Distance in latitude from the equator** | **-1.02 [-1.75, -0.32]** | **49151** | **0.005**** |
| | | Phylogenetic variance | 33.12 [0.0002, 153.34] | 8929 | - |
| | | Species | 364.06 [116.51, 714.05] | 31124 | - |
| | | Residual variance | 131.67 [73.94, 201.51] | 48733 | - |
| Fig 5B | B. CA at individual level in the opaque zone | Intercept | 158.67 [153.40, 164.90] | 3500 | 0.0003*** |
| | | Distance in latitude from the equator | -0.10 [-0.45, 0.25] | 3500 | 0.57 |
| | | Phylogenetic variance | 12.71 [0.0003, 63.41] | 3500 | - |
| | | Species | 16.37 [0.0002, 56.25] | 3948 | - |
| | | Residual variance | 46.97 [20.74, 78.05] | 3500 | - |
| | Question: Does transparency cover a lower wing surface area in tropical species compared to temperate species? Full model : Proportion of wing surface area covered by transparent patches ~distance in latitude from the equator 2 random factors tested: Phylogeny, Species | | | | |
| | C. Proportion of transparency on the wing at individual level in the transparent zone | Intercept | 39.24 [8.74, 69.83] | 3325 | 0.013* |
| | | Distance in latitude from the equator | -0.13 [-0.74, 0.47] | 3500 | 0.669 |
| | | Phylogenetic variance | 1160.59 [532.46, 1958.04] | 3500 | - |
| | | Residual variance | 20.30 [11.38, 31.18] | 3500 | - |
